# Top predators in soil food webs increase carbon cycling efficiency

**DOI:** 10.64898/2026.07.20.739503

**Authors:** Justine D. M. Lejoly, Esther van Hoof, Yuxin Wang, Valentin Favre, Casper W. Quist, Stefan Geisen, G.F. (Ciska) Veen

## Abstract

Soil microbes are considered central in litter decomposition and soil carbon formation. Microbial activity and abundance are controlled by microbivores, which are themselves preyed upon by top predators. However, the role of top predators in carbon cycling is rarely studied, especially in bacterivore-dominated food webs. Here we tested how trophic cascades, consisting of microbes, microbivores (bacterivore-dominated nematode communities) and top predators (nematode-feeding mites) impact carbon cycling and associated microbial pools and processes. We found that our model top predator decreased the abundance of fungivorous nematodes and had cascading effects on microbiome composition, notably increasing Gram-positive bacterial biomass, thus promoting the bacterial energy channel. These trophic cascades propagated to carbon cycling, decreasing heterotrophic respiration by 10 % while maintaining litter decomposition rates. Taken together, our results suggest that top predators increase carbon cycling efficiency and highlight the importance of complex trophic interactions, including trophic cascades, in determining soil carbon cycling.

## Introduction

Predators play a key role in driving ecosystem functioning, including biogeochemical cycles (Otto et al., 2008; Rizzuto et al., 2024; Schmitz et al., 2018). Their impact primarily takes place via trophic cascades, that is by downward propagation of top-down control across multiple trophic levels (Ripple et al., 2016). In aboveground food webs, when preying on herbivores, predators can indirectly increase primary productivity by reducing herbivory pressure, with positive consequences for carbon (C) storage (Otto et al., 2008; Strickland et al., 2013). Beyond controlling prey biomass, predators can also affect their metabolism and behavior, which in turn impacts biogeochemical cycling (Schmitz et al., 1997; Sommer et al., 2023). In soil food webs, top predators (sensu Thakur and Geisen, 2019) can control microbivore abundance (Laakso and Setälä, 1999), with potential cascading effects on microbiome composition and functioning (Thakur and Geisen, 2019), as well as C cycling, but experimental evidence is scarce (Lejoly et al., 2026).

Soil microbes (e.g., bacteria and fungi) contribute to soil organic matter (SOM) formation notably through litter decomposition and transformations into biomass and other microbial products with the help of extracellular enzymes (Cotrufo et al., 2013; Kallenbach et al., 2016). Microbial residues can further accumulate as necromass, an important precursor of stable SOM (Liang et al., 2019). The lower the proportion of mineralized C (measured as heterotrophic respiration) relative to the C transformed into microbial products, the higher the microbial efficiency to transform litter and other organic inputs into persistent SOM (Kallenbach et al., 2016; Mielke et al., 2022). This microbial C efficiency is largely determined by microbial community composition and resource availability. For example, higher relative fungal abundance is generally associated with higher microbial efficiency and SOM formation (Kallenbach et al., 2016). In addition, fast-growing Gram(-) bacteria primarily utilize labile C sources while slow-growing Gram(+) bacteria prefer more complex C compounds (Fanin et al., 2019).

There is growing evidence that microbivores, such as nematodes and protists, can directly affect microbial C efficiency by altering microbial biomass, activity, and community composition (Crowther et al., 2011; Kane et al., 2022; Trap et al., 2016), and therefore their impact on C cycling. Previous research has shown that microbivores decrease microbial biomass while increasing heterotrophic respiration, leading to a decrease in C cycling efficiency (Geisen et al., 2021; Trap et al., 2016). However, how higher trophic levels, including top predators, can further modulate these pathways and influence soil microbes and C cycling is unclear. The few manipulative experiments in the literature have mainly focused on fungal-dominated soil food webs (Chomel et al., 2019; Cortet et al., 2003; Hedlund and Öhrn, 2000; Lang et al., 2014; Lucas et al., 2020) while bacterial-dominated soil food webs are rarely studied in this context (but see Martikainen and Huhta, 1990; Zaitsev et al., 2018). There is however evidence that bacteria are more sensitive to top-down controls by higher trophic levels (Hunt et al., 1987; Martinez-Almoyna et al., 2022; Wardle and Yeates, 1993), suggesting that top predators may have stronger impact on C cycling via trophic cascades in bacterial than fungal-dominated food webs.

Here we tested the existence of trophic cascades in soils and consequences for litter decomposition and soil C cycling in grassland soil mesocosms by reconstructing bacterial-dominated soil food webs of increasing trophic complexity. We manipulated the presence of bacterivorous-dominated nematode communities and top predators in this model system (predatory mites) in a full factorial design. We investigated the response of the microbiome composition and biomass, of nematode abundance and feeding groups, and of C-cycling parameters (litter mass loss, heterotrophic respiration, potential hydrolytic extracellular enzyme activities, and microbial necromass) to the presence of top predators during five weeks of litter decomposition. We hypothesized that predatory mites would (1) decrease nematode abundance, (2) increase microbial biomass and necromass as well as alter microbiome composition, particularly increasing fast-growing Gram(-) bacteria by alleviating microbivory pressure, (3) increase C cycling efficiency by increasing litter decomposition rates while maintaining similar levels of heterotrophic respiration. We expected these effects to strengthen over the course of the experiment.

## Methods

### Soil and biota collection

Soil and litter were collected from a grassland on sandy soil (52°03’38”N 5°45’11”E; de Mossel, Veluwe, Ede, the Netherlands). Leaf litter of the three most common grass species (*Festuca rubra, Agrostis capillaris, Holcus lanatus*) was collected immediately upon senescence in October 2016. Soil was sampled in August 2022 from the same location from 0 – 10 cm depth and has the following properties: CaCl2-pH of 5.2, 3.2 % SOM, and 2 % Clay (Quist et al., 2019). Both soil and leaf litter were air-dried and sterilized by gamma radiation (> 25 KGray, isotron, Ede, the Netherlands) soon after collection.

In January 2023, fresh soil from 0 – 10 cm depth was collected for the extraction of microbial and nematode communities at the same location, for a total of six kg. Microbial communities were extracted in water (Hu et al., 2022; Li et al., 2020). In brief, 720 ml of demineralized water was added to 800 g of fresh soil. After stirring for 30 s and resting for 30 s, the supernatant was filtered through a 1 mm and then a 20 μm sieve, ensuring that nematodes were excluded (Li et al., 2020; Wang et al., 2019). The procedure was repeated to obtain sufficient volume for the inoculation of all mesocosms. Soil nematodes were extracted in batches of 500 g of fresh soil using an Oostenbrink elutriator and an additional cleaning step of three days with cotton wool filters (Oostenbrink, 1960) for a total of five kg. The extract was subsequently concentrated to obtain a water solution containing about 200 nematodes per ml. Initial identification confirmed that nematode communities were dominated by bacterivores.

Specimens of the predatory mite *Gaeolaelaps aculeifer* (G.Canestrini, 1884) were obtained from a laboratory culture.

### Experimental design and mesocosm setup

Mesocosms were set up with four different soil biota treatments: (1) three trophic levels (microbiome + nematodes + predatory mites, thereafter ‘with mites’ treatments), (2) two trophic levels (microbiome + nematodes, thereafter ‘without mites’ treatment), (3) one trophic level (microbiome, thereafter ‘microbial control’ treatment), and (4) non-trophic control (microbiome + predatory mites; Suppl. Fig. 1). Each treatment had eight replicates for each of the three destructive samplings (after one, two, and five weeks), which yields a total of 24 units per treatment and an additional eight mesocosms to be destructively harvested at the end of the pre-incubation period (4 soil treatments x 8 replicates x 3 harvests + 8 mesocosms for initial conditions = 104 experimental units). Each mesocosm consisted of 200 g of air-dried sterilized soil homogeneously mixed with 1.2 g of sterilized mixed grass litter (1:1:1 mass ratio of *Festuca rubra, Agrostis capillaris, Holcus lanatus*; C:N ratio of the mix: 17.5) and re-inoculated with 30 ml of the < 20 μm microbiome extract solution. An additional 0.3 g of mixed litter was added in a litter bag to monitor litter decomposition. To ensure sufficient re-colonization of the soil, all mesocosms were pre-incubated for three weeks in the dark with a temperature of 20 °C for 16 h and 15 °C for 8 h to simulate a day/night cycle. The establishment of the microbiome was checked by measuring heterotrophic respiration before soil food web reconstruction.

Once respiration rates had stabilized, three weeks after microbial inoculation, 48 mesocosms were inoculated with 10 ml of nematode extract (concentration of 200 ind. ml^-1^) to reach a concentration of 10 ind. g of soil^-1^, which is similar to natural densities observed at the site (Quist et al., 2019) and to the densities used in other experiments (Mielke et al., 2022). As the nematodes were in a water solution, the nematode-free mesocosms (48 + 8 mesocosms for initial conditions) received an equivalent volume (10 ml) of supernatant nematode-free water from the nematode solution to minimize nutrient and moisture differences.

A total of ten adult predatory mites were added to half of the mesocosms containing nematodes and half of the mesocosms containing only microbes (n=48), three days after the inoculation of nematodes to give the latter time to adapt to their new environment (Lang et al., 2014). The chosen density was similar to natural densities of predatory mites previously observed at the sampling site (1250 ind.m^-2^; de Groot et al., 2016). Eight mesocosms were destructively sampled at the time of the mite addition to determine initial conditions prior to the addition of fauna (Suppl. Table 1). Mesocosms were incubated in the dark with a temperature of 20 °C for 16 h and 15 °C for 8 h to simulate a day/night cycle and water content was kept at 20 %. Mesocosms were destructively sampled after one, two, and five weeks of incubation. For each harvest, after removal of the litter bag, the soil was homogenized and divided into subsamples for appropriate storage prior to chemical and biological analyses: freeze-dried for PLFA and microbial necromass, frozen for 16S rRNA gene sequencing and potential extracellular enzyme activities, and short-term fridge storage for microbial biomass and nematode extraction.

### Carbon fluxes and pools

Throughout the incubation, heterotrophic respiration was periodically measured by closing the mesocosm for 2 h after flushing with CO_2_-free air (Linde Gas, Schiedam, the Netherlands) for 2 min and then sampling 12 ml of headspace air into pre-vacuumed Exetainer vials, prior to measurement on a CH_4_/CO_2_ analyzer GC-Trace1600-TCD (Thermo Scientific, Bleiswijk, the Netherlands). Cumulative heterotrophic respiration was estimated by integration between the different measurements, considering a linear evolution of respiration rates and using the ideal gas law. It was assumed that heterotrophic respiration was of microbial origin, as faunal respiration is negligible (Mikola and Setälä, 1998).

Litter decomposition was estimated using a nylon mesh bag of 5 cm × 10 cm and mesh size 0.9 × 1 mm filled with 0.3 g of sterilized mixed grass litter (1:1:1 mass ratio of *Festuca rubra, Agrostis capillaris, Holcus lanatus*) placed under the soil surface. Litter mass loss was calculated by subtracting the remaining litter mass from the initial litter mass after loss-on-ignition for 3 h at 550 °C, as ash-free mass loss, after one, two, and five weeks of experiment and accounting for initial losses from manipulation by including traveler bags.

Microbial biomass C and N (MBC and MBN) were estimated by the chloroform-fumigation-extraction method as the difference in dissolved organic C (DOC) and dissolved N (DN) between fumigated and non-fumigated fresh soil samples (15 g) after one, two, and five weeks of experiment, as well as for initial conditions (Vance et al., 1987), using a Shimadzu TOC-L analyser coupled with a TNM-L unit (Shimadzu Benelux, ‘s-Hertogenbosch, the Netherlands).

Combining litter decomposition and heterotrophic respiration allowed us to estimate C cycling efficiency. Slower litter decomposition rates combined with higher or similar heterotrophic respiration would indicate lower C cycling efficiency, while lower heterotrophic respiration combined with similar or higher litter decomposition rates would indicate higher C cycling efficiency.

### Microbial necromass

Amino sugars were extracted from freeze-dried samples to estimate microbial necromass after five weeks of experiment, as well as for the initial conditions (Joergensen, 2018; Salas et al., 2023). In brief, 0.5 g of freeze-dried soil was hydrolyzed for 6 h at 105 °C with 6M HCl. After centrifugation, the supernatant was separated and processed according to Salas et al. (2023). Derivatized samples were measured on an ultra-high performance liquid chromatograph – mass spectrometer (UHPLC-MS) 1290 Infinity II connected to a 6490C triple quadruple MS (Agilent, Santa Clara, USA). The compounds were separated on an ACCQ-TAG ULTRA C18 column (1.7 um, 2.1×150 mm, Waters, Milford, USA) and quantified using external standards. As our method was unsuccessful in separating glucosamine from mannosamine, we calculated the sum of all four amino sugars (glucosamine, mannosamine, muramic acid, galactosamine) as a proxy for total microbial necromass (Liang et al., 2013). Two samples were removed from the dataset as the concentration was lower than the limit of detection, showing that the extraction was likely unsuccessful (Indorf et al., 2011).

### Phospholipid fatty acid analysis

Phospholipid fatty acid (PLFA) analysis was used to estimate the abundance of different microbial groups after one and five weeks, as well as for initial conditions. In brief, polar lipids were extracted from 2 g of freeze-dried soil using a Bligh and Dyer solution (chloroform:methanol:citrate buffer 1:2:0.8 v/v/v). Extracted phospholipids were separated using solid-phase extraction columns (SPE Bond Elut 1cc LRC-SI, 100 mg) and derivatized into fatty acid methyl esters (FAMEs) using mild alkaline trans methylation. FAMEs were then analyzed on an Agilent 7890A gas chromatograph (Agilent Technologies, Wilmington, USA) coupled to a flame ionization detector and equipped with an HP-5MS 60 m column. The internal standard methyl nonadecanoate (19:0) was used to calculate PLFA concentrations and compounds were identified using an Equivalent Chain Length compound list for common fatty acids together with standard bacterial fatty acid methyl ester and FAME mixes. Specific PLFAs were used as markers of the following microbial groups: 18:2ω6 for fungi, the sum of i14:0, i15:0, a15:0, i17:0, and a17:0 for Gram(+) bacteria, and the sum of cy17:0, cy19:0, 16:1ω7, and 18:1ω9 for Gram(-) bacteria (Orwin et al., 2018; Saetre, 1998; Stromberger et al., 2012; Zelles, 1999). The fungi:bacteria ratio was calculated by dividing 18:2ω6 by the sum of all Gram(+) and Gram(-) bacteria markers.

### 16S rRNA gene sequencing

Bacterial community composition was characterized using 16S rRNA gene sequencing at the end of the experiment (after five weeks). In brief, DNA was extracted from 1 g of thawed soil following Harkes et al. (2025) and sent to Quebec Genome (Quebec, Canada) for sequencing of the V4 region of the 16S ribosomal RNA gene using the primers 515F-Y (5’-GTGYCAGCMGCCGCGGTAA-3’) and 926R (5’-CCGYCAATTYMTTTRAGTTT-3’) (Parada et al., 2016). This resulted in a total of 726,058 reads across all samples, with a minimum of 10,867 reads per sample. The demultiplexed raw reads were filtered and denoised with DADA2 using QIIME2 (Callahan et al., 2016; Quast et al., 2013). Both forward and reverse primers were trimmed, and only reads with fewer than five expected errors maxEE = 5) were retained (Edgar and Flyvbjerg, 2015). Chimeric sequences were removed, and the remaining high-quality sequences were used to generate Amplicon Sequence Variants (ASVs) based on the SILVA v138.1 database (Quast et al., 2013). Taxonomic classification was performed using the scikit-learn method (Pedregosa et al., 2011) with a Naive Bayes classifier trained specifically for our primer pairs, using the Reference Sequence Annotation and Curation Pipeline (Li et al., 2021). The taxonomic information obtained was combined with ASV abundance data for statistical analysis.

### Potential extracellular enzyme activities

Potential hydrolytic extracellular enzyme activity assays were conducted for C-cycling (β-1,4-glucosidase, cellobiohydrolase, and β-1,4-xylosidase), N-cycling (β-1,4-N-acetyl-glucosaminidase and leucine aminopeptidase), P-cycling (acid phosphatase), and S-cycling (arylsulfatase) enzymes following an adapted method from Baldrian (2009) using 1 g of thawed soil. Potential enzyme activities were estimated by measuring fluorescence after a 2-hour incubation with corresponding fluorogenic substrates in the dark at 40 °C, using a BioTek Synergy HTX Multimode Reader with an excitation wavelength of 360 nm and an emission wavelength of 460 nm (Beun De Ronde, Abcoude, the Netherlands). The fluorogenic substrates used contained 4-methylumbelliferyl for all enzymes, except leucine aminopeptidase for which 4-methyl coumarin was used. Concentrations were calculated using standard curves, after correction for background fluorescence of soil and substrates. The sum of these seven hydrolytic extracellular enzyme activities was used as an estimate of total potential hydrolytic enzyme activity.

### Nematode abundance and feeding groups

After each destructive sampling, nematodes were extracted from 70 g of fresh soil with an Oostenbrink elutriator. Nematodes were counted and classified into feeding groups (herbivore, bacterivore, fungivore, and carnivore) based on mouth part morphology (Yeates et al., 1993) using a reverse light stereo microscope Leica DMIL (Leica Microsystems GmbH, Wetzlar, Germany). The channel index was calculated as the ratio between fungivorous and bacterivorous nematodes.

### Statistical analyses

All statistical analyses were performed in R version 4.3.2 (R Core Team, 2023) with a focus on the last destructive sampling (week 5). Potential impacts of soil food web composition on nematodes (feeding group abundances) and microbes (Gram(+) and Gram(-) bacteria, and fungi), as well as on C-cycling related parameters (heterotrophic respiration, litter mass loss, microbial biomass C and N, amino sugars, extracellular enzyme activities) were tested using one-way analysis of variance with biota treatment as factor, followed by Tukey posthoc test from the agricolae R package (de Mendiburu, 2020). Data was transformed using Tukey’s ladder of power to reach normality of residuals (Mangiafico, 2021) for leucine aminopeptidase, cellobiohydrolase, β-1,4-glucosidase, acid phosphatase, muramic acid, glucosamine + mannosamine, and fungi. When the assumption of homogeneity of variance was violated, non-parametric test Kruskal-Wallis was used instead of one-way analysis of variance (for β-1,4-N-acetyl-glucosaminidase after two weeks, as well as total and bacterivorous nematode abundance after one week). Trophic relationships within nematode communities were investigated using linear regressions with lm function, through stat_poly_eq embedded in ggplot. The threshold for statistical significance was set at *p* < 0.05.

For a subset of parameters (PLFA, microbial biomass C and N, litter mass loss, and extracellular enzyme activities), additional statistical analyses were conducted for the first two destructive sampling (after one and two weeks of experiment) and showed that most parameters were not yet affected by the biota treatments (Suppl. Fig. 2-5; Suppl. Tables 2-3).

16S rRNA gene amplicon sequencing data were analyzed using the microeco R package (Liu et al., 2021). ASVs were rarefied to 10,500 reads per sample to account for differences in sequencing depth before downstream statistical analysis. Differences in bacterial community composition were determined using permutational analysis of variance with Bray-Curtis distance and visualized by principal coordinates analysis. Changes in relative abundance of phyla accounting for > 5 % relative abundance and in alpha-diversity (Shannon index) were further investigated using one-way analysis of variance as described above.

## Results

### Nematode communities

After one week, the presence of predatory mites did not affect nematode feeding group abundances (Suppl. Fig. 2 & Suppl. Table 2) but after two weeks, the total abundance of nematodes as well as that of bacterivorous and fungivorous nematodes decreased when predatory mites were present (Suppl. Fig. 2 & Suppl. Table 3). After five weeks, the presence of predatory mites resulted in a 50% decrease in fungivorous (*F*_1,14_ = 7.7, *p* < .05; Fig. 1C) and a 75% decrease in carnivorous nematode abundances (*F*_1,14_ = 20.2, *p* < .001; Fig. 1D) but did not affect bacterivorous nor total nematode abundance (Fig. 1A-B). Mites also affected prey-predator relationships within nematode communities. Fungivore abundance was negatively correlated with carnivore abundance only in the presence of predatory mites (Fig. 1E), while bacterivore abundance was not correlated with carnivore abundance in any treatment (Fig. 1F). However, predatory mites decreased the ratio of fungivorous-to-bacterivorous nematodes (*F*_1,14_ = 5.5, *p* < .05; Fig. 2B).

**Figure 1.**
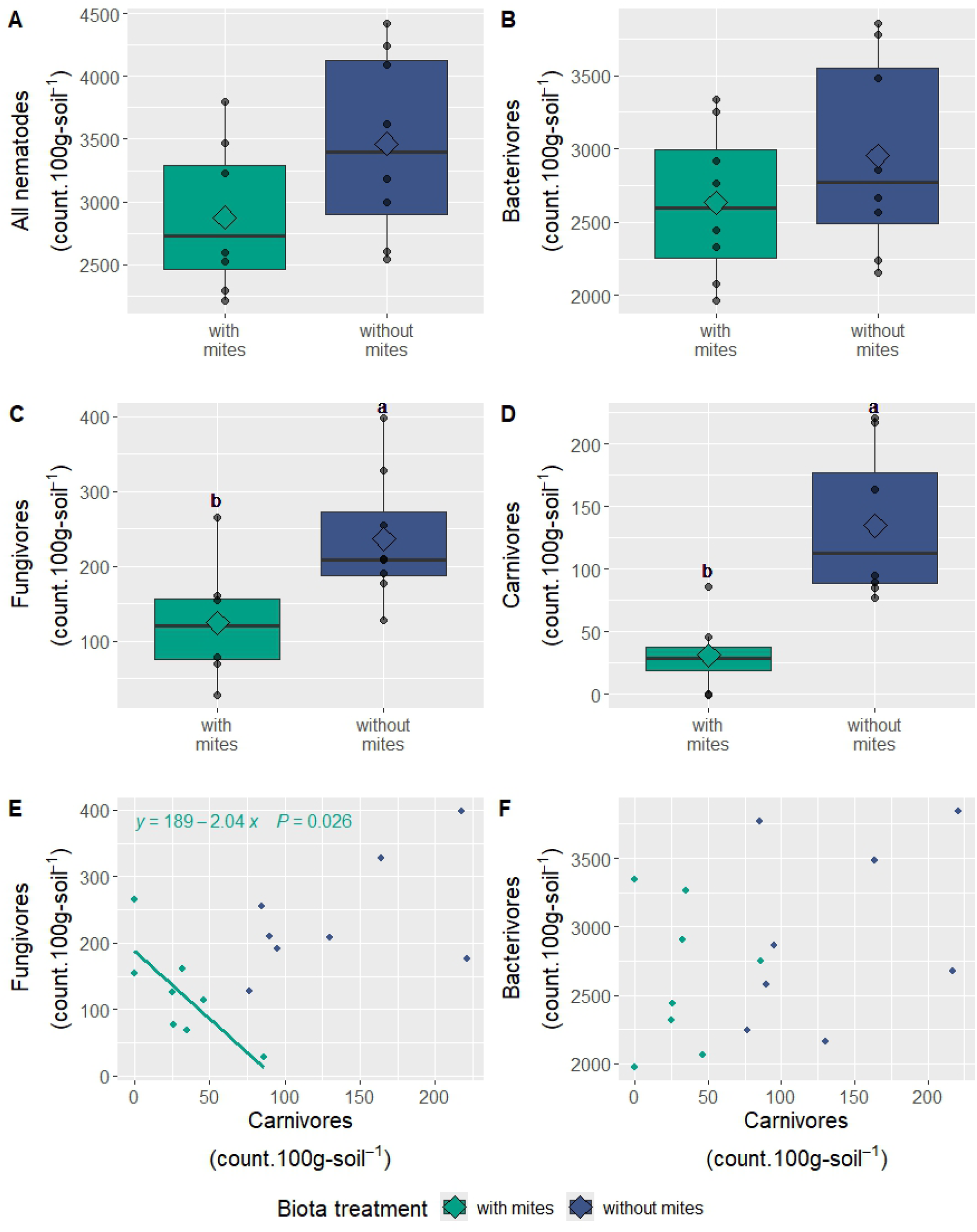
Response of nematode communities to mite presence. Median and 25^th^ and 75^th^ percentiles (corresponding to the lower and upper hinges) of abundances of all (A), bacterivorous (B), fungivorous (C), and carnivorous (D) nematodes, as well as the trophic relationships between carnivorous nematodes and their preys: fungivorous (E) and bacterivorous (F) nematodes after five weeks with different biota treatments (with mites: microbiome (M) + nematodes (N) + predatory mites (P), without mites: M + N, n = 8, only treatments containing nematodes are displayed in the figure). The mean and individual data points are depicted as diamond and dots, respectively. Different letters indicate significant differences between treatments, according to one-way analysis of variance. Absence of letters means absence of significant differences (p > .05) for the pairwise posthoc Tukey HSD test. The upper whisker extends from the hinge to the largest value no further than 1.5 * the inter-quartile range (IQR) from the hinge. The lower whisker extends from the hinge to the smallest value at most 1.5 * IQR of the hinge. Data beyond the end of the whiskers are considered outliers. For the regressions, only significant relationships are shown on the graph, including equation and associated p value.

**Figure 2.**
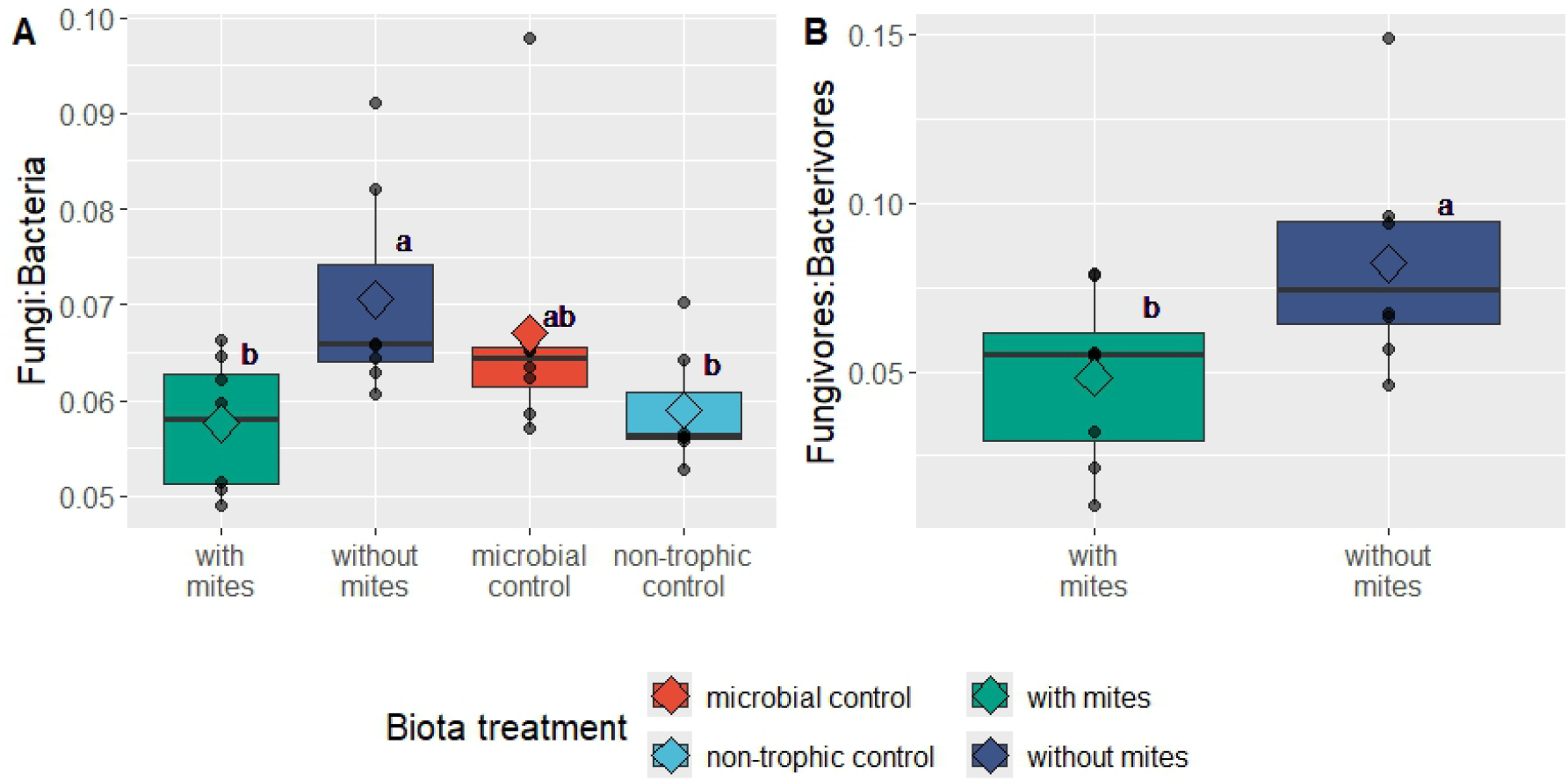
Relative prevalence of bacterial versus fungal energy channels. Median and 25^th^ and 75^th^ percentiles (corresponding to the lower and upper hinges) of ratios of fungi:bacteria biomass (A) and fungivorous-to-bacterivorous nematodes (B) after five weeks (n = 8) with different biota treatments (with mites: microbiome (M) + nematodes (N) + predatory mites (P), without mites: M + N, microbial control: M, non-trophic control: M + P). The mean and individual data points are depicted as diamond and dots, respectively. Different letters indicate significant differences between treatments, according to one-way analysis of variance. Absence of letters means absence of significant differences (p > .05) for the pairwise posthoc Tukey HSD test. The upper whisker extends from the hinge to the largest value no further than 1.5 * the inter-quartile range (IQR) from the hinge. The lower whisker extends from the hinge to the smallest value at most 1.5 * IQR of the hinge. Data beyond the end of the whiskers are considered outliers.

### Microbial communities

After five weeks, Gram(+) bacterial biomass was higher with mites compared to all other treatments (*F*_3,28_ = 3.5, p < .05; Fig. 3A). This trend was already partly visible after one week, with higher biomass with mites compared to the microbial control (Suppl. Fig. 3 & Suppl. Table 2). Fungal and Gram(-) bacterial biomasses were not affected by the biota treatments after one (Supp. Fig. 3 & Suppl. Table 2) and five weeks (*F*_3,28_ = 2.4, *p* = .088 and *F*_3,28_ = 0.3, *p* = .829, respectively; Fig. 3B-C). However, the fungi:bacteria ratio was lower with than without mites (Fig. 2A).

**Figure 3.**
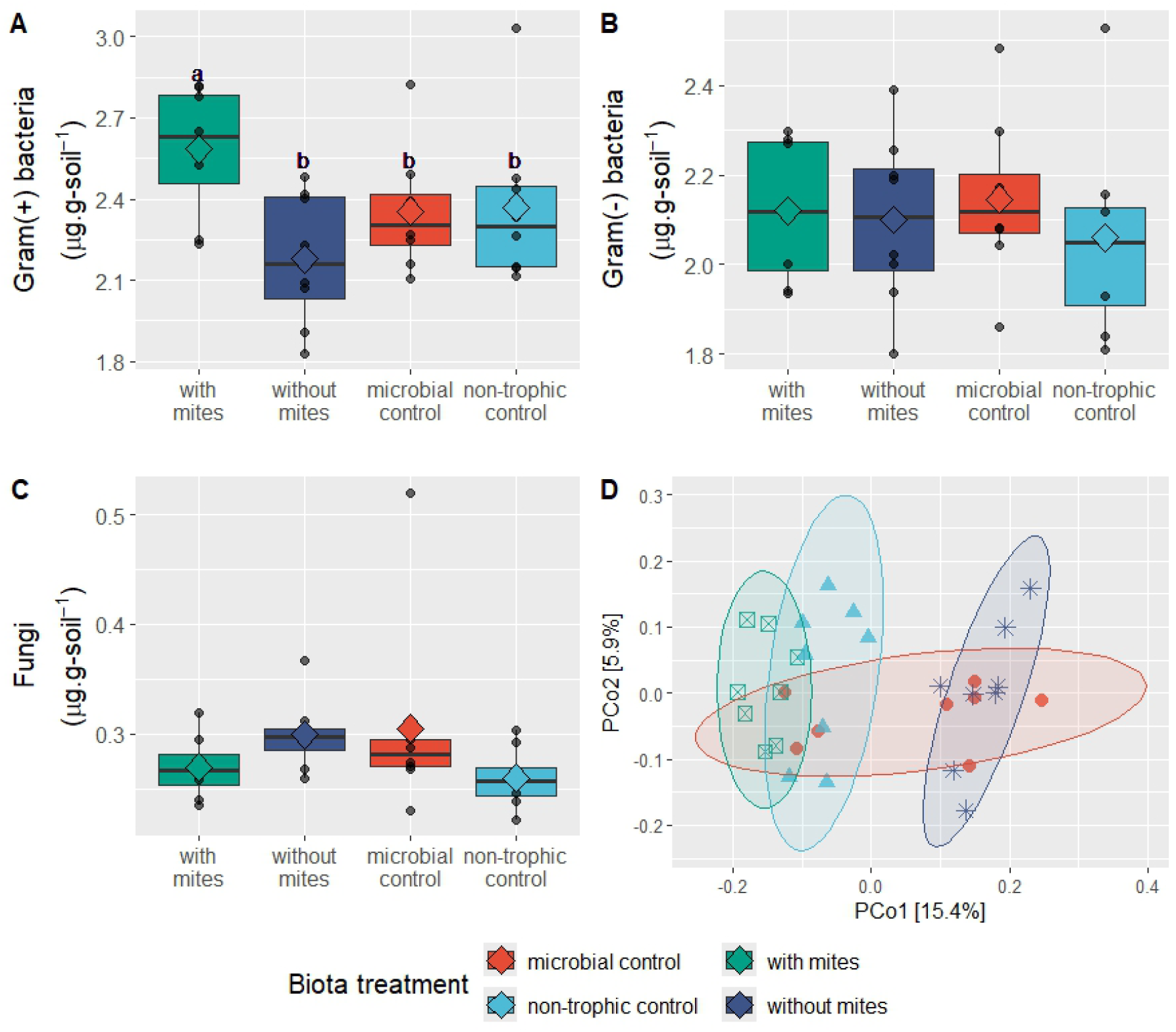
Response of microbial communities to soil biota treatments. Median and 25^th^ and 75^th^ percentiles (corresponding to the lower and upper hinges) of biomasses of (A) Gram(+) bacteria, (B) Gram(-) bacteria, and (C) fungi after five weeks with different biota treatments (with mites: microbiome (M) + nematodes (N) + predatory mites (P), without mites: M + N, microbial control: M, non-trophic control: M + P), as well as the mean and individual data points depicted as diamond and dots, respectively, from the analysis of phospholipid fatty acids (n=8). Different letters indicate significant differences between treatments, according to one-way analysis of variance. Absence of letters means absence of significant differences (p > .05) for the pairwise posthoc Tukey HSD test. The upper whisker extends from the hinge to the largest value no further than 1.5 * the inter-quartile range (IQR) from the hinge. The lower whisker extends from the hinge to the smallest value at most 1.5 * IQR of the hinge. Data beyond the end of the whiskers are considered outliers. The principal coordinate analysis of 16S rRNA gene amplicon sequencing data using Bray-Curtis distance is displayed in (D).

Bacterial community composition differed between biota treatments, with the treatments with and without mites being the most different from each other (Fig. 3D & Suppl. Table 1; *pseudo-F*_3,28_ = 2.23, *p* < .001). In contrast, Shannon α-diversity was not affected by food web composition (Suppl. Fig. 4). The relative abundance of Actinobacteriota increased by 68 % with compared to without mites while the opposite pattern was observed for Proteobacteria, decreasing by 21 % (Suppl. Fig. 5 & Suppl. Table. 2).

After one and two weeks, none of the potential extracellular enzyme activities were affected by the biota treatments (Suppl. Table 2-3; Suppl. Fig. 6-7). At the end of the experiment, after five weeks, only leucine aminopeptidase and β-1,4-N-acetyl-glucosaminidase were affected by the biota treatments (Table 1 & Suppl. Table 4). Pairwise comparisons revealed that leucine aminopeptidase activity was significantly higher in the non-trophic control than in the treatment without mites, while differences in β-1,4-N-acetyl-glucosaminidase activity were non-significant according to the posthoc test.

**Table 1.** Averages and standard errors for microbiome biomass, necromass, and extracellular enzyme activities after five weeks with different biota treatments (with mites: microbiome (M) + nematodes (N) + predatory mites (P), without mites: M + N, microbial control: M, non-trophic control: M + P). Different letters indicate significant differences based on one-way analysis of variance with p < .05 (n=8 except for microbial necromass where n=7 for ‘with mites’ and ‘without mites’ treatments).

|  | With mites | Without mites | Microbial control | Non-trophic control |
| --- | --- | --- | --- | --- |
| **Microbial biomass nitrogen ($\mu\text{g.g-soil}^{-1}$)** | 0.5 (0.2) a | 0.7 (0.2) a | 0.9 (0.2) a | 0.8 (0.2) a |
| **Microbial necromass ($\mu\text{M.l}^{-1}$)** |  |  |  |  |
| Muramic acid | 3.7 (0.6) a | 2.9 (1.3) a | 4.2 (0.9) a | 2.5 (0.7) a |
| Glucosamine + Mannosamine | 12.3 (1.8) a | 9.9 (2.0) a | 13.8 (2.0) a | 10.9 (1.6) a |
| Total microbial necromass | 70.4 (6.9) a | 51.9 (5.6) a | 58.9 (6.5) a | 63.1 (5.6) a |
| **Extracellular enzyme activities ($\text{nmol.g-soil}^{-1}.\text{h}^{-1}$)** |  |  |  |  |
| Leucine aminopeptidase | 33.1 (3.0) ab | 21.8 (2.6) b | 27.3 (2.3) ab | 39.7 (7.4) a |
| Cellobiohydrolase | 81.3 (14.7) a | 89.3 (9.9) a | 107.0 (4.0) a | 93.64 (5.7) a |
| β-1,4-glucosidase | 286.9 (34.7) a | 319.5 (37.1) a | 387.8 (57.6) a | 392.4 (47.7) a |
| β-1,4-N-acetyl-glucosaminidase | 76.1 (7.6) a | 80.8 (8.8) a | 99.3 (5.9) a | 100.7 (6.1) a |
| Acid phosphatase | 179.4 (27.1) a | 269.0 (70.4) a | 227.5 (27.8) a | 241.3 (43.5) a |
| Arylsulfatase | 62.6 (6.3) a | 66.7 (7.8) a | 82.4 (7.6) a | 80.2 (6.7) a |
| β-1,4-xylosidase | 67.3 (8.4) a | 75.7 (7.5) a | 89.8 (7.0) a | 90.6 (8.9) a |
| Total potential extracellular enzyme activity | 1021.1 (80.0) a | 922.6 (78.9) a | 786.8 (69.6) a | 1038.4 (67.2) a |

### Carbon fluxes and pools

The absence of predatory mites increased cumulative CO_2_ respiration by 10 % over the five-week incubation (*F*_3,28_ = 10.2, *p* < .001; Fig. 4A). After one week, litter mass loss was higher with than without mites (Suppl. Fig. 8A; Suppl. Table 2), but differences disappeared after two (Suppl. Fig. 8B; Suppl. Table 3) and five weeks (*F*_3,28_ = 0.3, *p* = .852; Fig. 4B). Total amino sugar concentrations, used as a proxy for microbial necromass, were not affected by the biota treatments (*F*_3,26_ = 1.3, *p* = .279; Table 1), but galactosamine concentrations were higher with than without mites, although this effect was not apparent in the posthoc test (*F*_3,26_ = 2.9, *p* = .052; Fig. 4C). Microbial biomass C and N were not significantly affected by the biota treatments after one, two (Supp. Fig. 9; Suppl. Table 2-3), and five weeks of experiment (Table 1; *F*_3,28_ = 1.9, p = .146; Fig. 4D).

**Figure 4.**
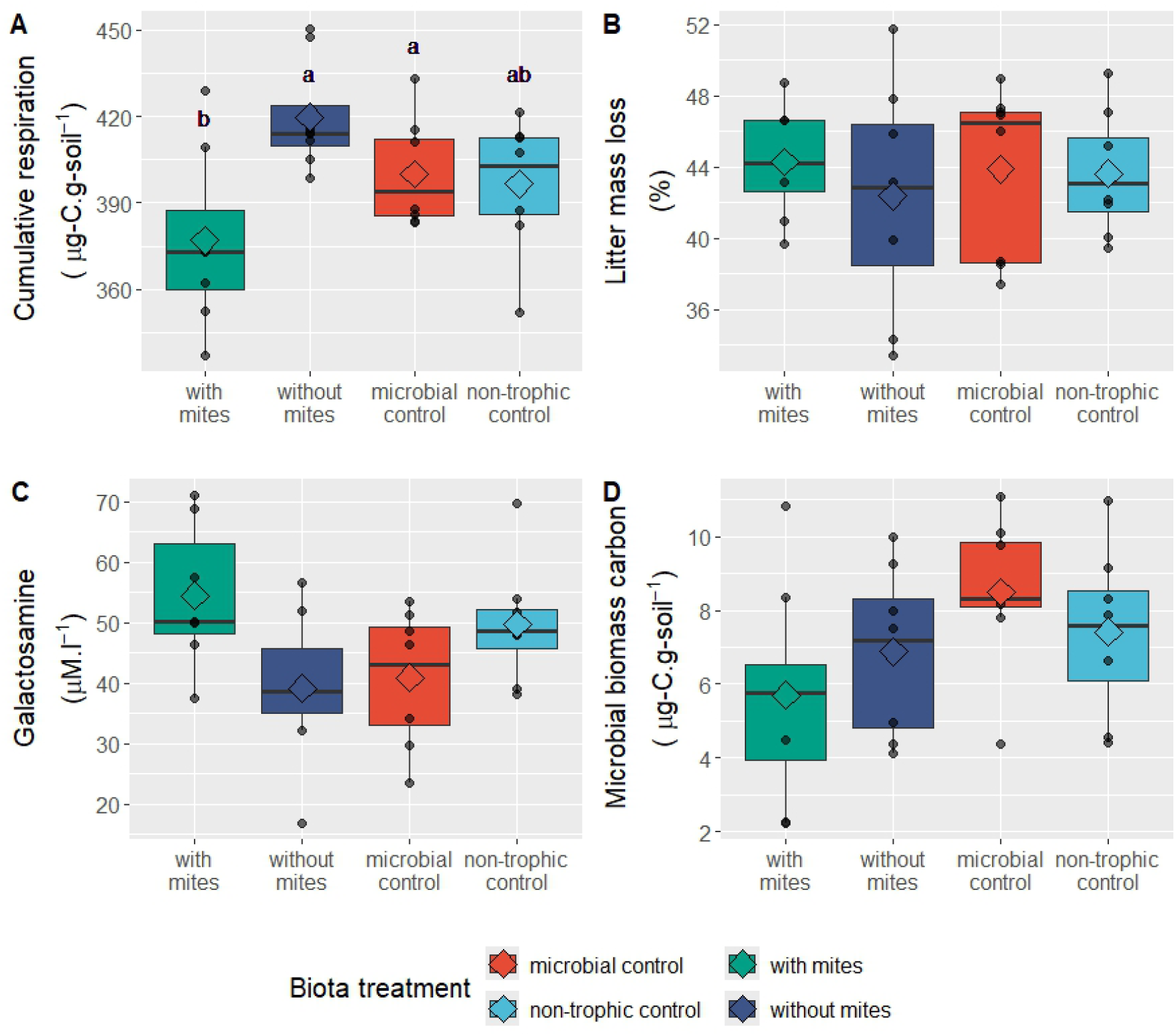
Response of carbon cycling parameters to soil biota treatments. Median and 25^th^ and 75^th^ percentiles (corresponding to the lower and upper hinges) of (A) cumulative respiration, (B) litter mass loss, (C) galactosamine, and (D) microbial biomass carbon after five weeks with different biota treatments (with mites: microbiome (M) + nematodes (N) + predatory mites (P), without mites: M + N, microbial control: M, non-trophic control: M + P), as well as the mean and individual data points depicted as diamond and dots, respectively (n=8, except for galactosamine where n=7 for the treatments ‘with mites’ and ‘without mites’). Different letters indicate significant differences between treatments, according to one-way analysis of variance. Absence of letters means absence of significant differences (p > .05) for the pairwise posthoc Tukey HSD test. The upper whisker extends from the hinge to the largest value no further than 1.5 * the inter-quartile range (IQR) from the hinge. The lower whisker extends from the hinge to the smallest value at most 1.5 * IQR of the hinge. Data beyond the end of the whiskers are considered outliers.

## Discussion

Our grassland soil mesocosm experiment with reconstructed bacterial-dominated soil food webs presents evidence of trophic cascades from top predators to C cycling, through changes in nematode and bacterial communities. Although such indirect trophic effects are largely ignored in biogeochemical conceptual frameworks and models (Angst et al., 2024; Lejoly et al., 2026), we showed that top predators can shift microbiome composition and alter C cycling. Combining multiple C cycling and associated biological (faunal and microbial) parameters enabled us to identify pathways through which these trophic cascades occur (Fig. 5).

**Figure 5.**
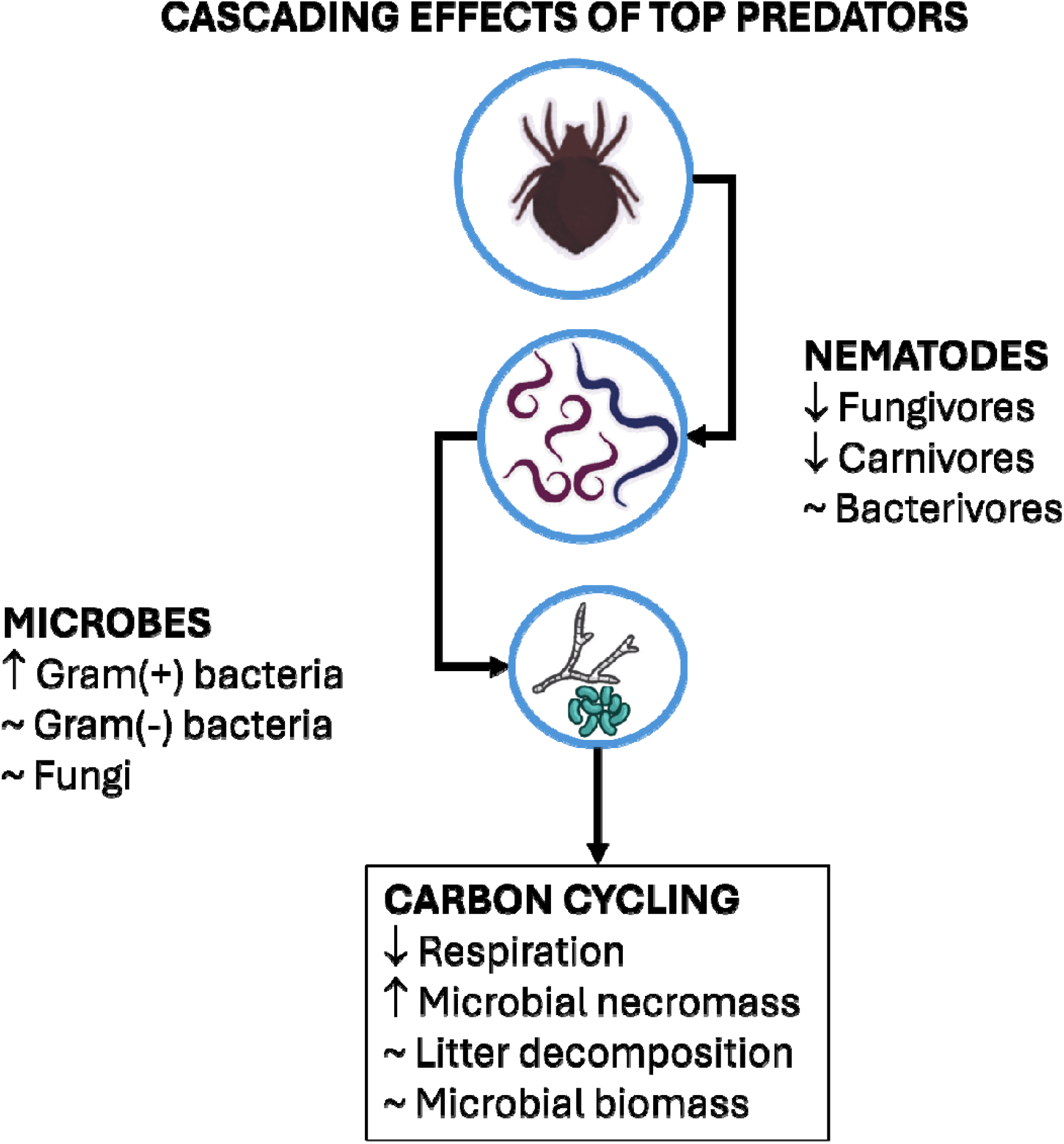
Conceptual figure depicting the pathways by which top predators (in this case, of predatory mites) influences nematode and microbial communities, with cascading effects on carbon (C) cycling. Top predators decrease fungivorous and carnivorous nematodes, which results in an increase in Gram(+) bacteria and higher C cycling efficiency, as respiration decreases and microbial necromass increases for similar litter decomposition rates. Changes in microbial and nematode communities indicate that predators promote the bacterial over the fungal energy channels.

### Top predators promote the bacterial energy channel by reducing fungivorous nematodes and promoting Gram(-) bacteria

Partly confirming our first hypothesis, we found that predatory mites in soil food webs decreased the abundance of fungivorous and carnivorous nematodes. However, the pathways were different from what we expected: top predators did not directly decrease the absolute abundance of bacterivorous nematodes, the dominant feeding group in our mesocosms, but instead increased their relative contribution to energy channels compared to fungivorous nematodes. While other studies have also reported a reduction in prey abundance (here, nematodes) with the presence of their predator (here, predatory mites) in reconstructed soil food webs (Chomel et al., 2019; Martikainen and Huhta, 1990) and in other ecosystem types (Shurin et al., 2002), changes in the relative contribution of bacterial and fungal energy channels are not typically presented.

Partly in line with our second hypothesis, we found that predatory mites altered microbiome composition but did not increase the biomass of Gram(-) bacteria. Instead, top predators increased the biomass of Gram(+) bacteria and the relative abundance of Gram(+) Actinobacteria, viewed as more resistant to disturbances (de Vries and Shade, 2013). Another study investigating the impacts of microbivory on microbial biomass, but without top predators, found Actinobacteria to be negatively impacted by both fungivores and bacterivores (Mielke et al., 2022). We now show that by adding top predators, microbivory pressure on microbial communities is lessened, with positive effects on Actinobacteria and a higher bacterial biomass without changes in fungal biomass. Taken together, our results show that top predators overall promoted the bacterial energy channel, with relatively higher contributions of both bacterivorous nematodes and bacteria. These energy channel shifts are likely to affect the overall resilience and resistance of the microbial communities, as well as their functioning (de Vries and Shade, 2013; Kallenbach et al., 2016; Prommer et al., 2020).

Reductions in fungivorous and carnivorous nematodes can be a result of two mechanisms: predatory mites may preferably feed on larger, more nutritious preys (e.g., larger carnivorous nematodes) and/or faster growing bacterivorous nematodes may have sufficient compensatory growth (Sechi et al., 2018). In our experimental setup, bacterivorous nematodes have two potential predators: predatory mites and carnivorous nematodes. Even though the abundance of bacterivorous nematodes was not affected by top predators, their metabolic rate could have been impacted (Hawlena and Schmitz, 2010; Schmitz et al., 2004). In the presence of top predators, the increase in Gram(+) bacteria, which are typically slow growers (de Vries and Shade, 2013), suggests lower bacterivory pressure, which could be in line with lower metabolic rates of bacterivorous nematodes associated with lower predation pressure. Taken together these results hint towards bacterivorous nematodes being primarily preyed upon by carnivorous nematodes rather than predatory mites, adding a fourth trophic level to this trophic cascade: predatory mites feed on carnivorous nematodes, which feed on bacterivorous nematodes, which in turn feed on bacteria.

### Top predators have limited impacts on microbial pools and enzyme activity

Contrary to our second hypothesis, we did not find any effects of top predators on microbial biomass nor on potential extracellular enzyme activity. Few studies have investigated the impact of microbivores and top predators on potential extracellular enzyme activity and all focused on fungal-dominated food webs (Crowther et al., 2011; Lucas et al., 2020). Our findings may indicate that top predators have less influence on potential extracellular enzyme activity in bacterial-dominated soil food webs or that the energetically costly production of enzymes was not needed as nutrient availability was sufficient throughout the experiment (Sinsabaugh et al., 2008). Sufficient nutrient availability could also explain why top predators did not increase microbial biomass, as microbes may be less sensitive to top-down control in the absence of nutrient limitations (Lenoir et al., 2007). However, we found evidence that top predators in soil food webs can partly promote the formation of microbial necromass, an important pool of stable SOM (Liang et al., 2019). The observed higher microbial necromass can be linked to the increase in Gram(+) bacteria, which have higher glucosamine concentrations than Gram(-) bacteria (Liang et al., 2019) and can also indicate higher microbial turnover in the presence of predators, as microbial biomass remained the same.

General conclusions cannot be drawn from the small number of studies investigating the role of top predators on microbial and carbon processes, especially considering the variety of experimental setups and analyses (Hedlund and Öhrn, 2000; Lucas et al., 2020; Martikainen and Huhta, 1990). Notably, previous research has identified that the type of resources available (humus, glucose, living plants) to microbes has a strong effect on the direction and strength of the impact of higher trophic levels (Lenoir et al., 2007; Lucas et al., 2020; Thakur et al., 2015). Multiple carbon sources should be considered in future experiments to investigate the role of top predators in different soil compartments (rhizosphere, detritusphere) or under different management scenarios (Lejoly et al., 2026). As most significant parameters were only affected at the end of the experiment, we expect that longer experiments could reveal the impact of trophic cascades on large SOM pools, such as mineral-associated and particulate organic matter (Lucas et al., 2020), and on later decomposition stages.

### Top predators promote carbon cycling efficiency by decreasing heterotrophic respiration while maintaining decomposition rates

Partly in line with our third hypothesis, we found that predators decreased cumulative heterotrophic respiration, but this was not associated with increased litter decomposition. The absence of change in litter decomposition matches previous studies (Martikainen and Huhta, 1990; Miyashita and Niwa, 2006), although Chomel et al. (2019) reported a decrease in the presence of top predators. Our study presents evidence that top predators can increase C cycling efficiency, by decreasing C losses while maintaining similar decomposition rates. In other words, for a similar amount of decomposed litter, microbes are mineralizing less C. Lucas et al. (2020) also found that predators increased microbial efficiency, by decreasing mass-specific respiration in both rhizosphere and bulk soil. Our study confirms that microbial efficiency is also higher in the early stages of litter decomposition (i.e., detritusphere) when top predators are present. The observed increase in Gram(+) bacteria resulting from trophic cascades may explain this higher C cycling efficiency, as these slow-growing bacteria are considered to have higher growth efficiency than fast-growing Gram(-) bacteria (Wieder et al., 2014).

Higher trophic levels such as top predators are generally more sensitive to global changes and disturbances (Lensing and Wise, 2006; Postma-Blaauw et al., 2010; Quist et al., 2020; Tsiafouli et al., 2015) and their loss may have consequences for soil C cycling. Since top predators promoted the bacterial energy channel in our study, predator loss could have the opposite effect, by promoting the fungal energy channel, considered less resilient to disturbances (de Vries and Caruso, 2016; Hedlund et al., 2004), and by negatively affecting bacteria resistant to disturbances such as Actinobacteria (de Vries and Shade, 2013). The cascading effects of top predators on soil C cycling may become more important with climate change (Lang et al., 2014). However, a previous meta-analysis indicated that biodiversity – C cycling relationships are stronger for low biodiversity, such as in our experimental setup with a single top predator taxon, and may disappear with increasing biodiversity (Nielsen et al., 2011). Future research should investigate whether similar mechanisms are also observed in soil food webs harboring higher predator diversity, how the relationships are influenced by predator density (Thakur et al., 2015), and whether the observed patterns persist in later decomposition stages.

## Acknowledgments

The predatory mites were provided by Diana Rueda-Ramirez. We thank Hans Zweers, Ciska Raaijmakers, Lauryn Padwick, Ivor Keesmaat, and Iris Chardon for their help with chemical analyses, and Joliese Teunissen for her contributions to bioinformatics analysis. J.D.M.L. acknowledges funding by the European Research Executive Agency for the Marie Skłodowska-Curie Actions Postdoctoral Fellowship Project no. 101105509 (Soil Fauna MIND). Views and opinions expressed are, however, those of the author(s) only and do not necessarily reflect those of the European Union or other funders. Neither the European Union nor the granting authority can be held responsible for them. G.F.V. acknowledges the Dutch Research Council (NWO) for funding of the Aspasia project 15.015.031. S.G. acknowledges the Deutsche Forschungsgemeinschaft (DFG, German Research Foundation) – Transregio Collaborative Research Centre 410 “WETSCAPES2.0”. Y.W. acknowledges China Scholarship Council (CSC) (grant no. 202104910024).

## Statement of authorship

JDML: conceptualization, data curation, formal analysis, investigation, methodology, funding acquisition, visualization, writing – original draft. EvH: investigation, writing – original draft, data curation. YW: investigation, data curation, writing – original draft. VF: investigation. CWQ: conceptualization, methodology, writing – review & editing. SG: conceptualization, funding acquisition, writing – review & editing. GFV: conceptualization, methodology, funding acquisition, supervision, writing – review & editing.

## References

Angst, G., Potapov, A., Joly, F.-X., Angst, Š., Frouz, J., Ganault, P., Eisenhauer, N., 2024. Conceptualizing soil fauna effects on labile and stabilized soil organic matter. Nat. Commun. 15, 1–16. 10.1038/s41467-024-49240-x

Baldrian, P., 2009. Microbial enzyme-catalyzed processes in soils and their analysis. Plant Soil Environ. 55, 370–378. 10.17221/134/2009-PSE

Callahan, B.J., McMurdie, P.J., Rosen, M.J., Han, A.W., Johnson, A.J.A., Holmes, S.P., 2016. DADA2: High resolution sample inference from Illumina amplicon data. Nat Methods 13, 581–583. 10.1038/nmeth.3869

Chomel, M., Lavallee, J.M., Alvarez-Segura, N., de Castro, F., Rhymes, J.M., Caruso, T., de Vries, F.T., Baggs, E.M., Emmerson, M.C., Bardgett, R.D., Johnson, D., 2019. Drought decreases incorporation of recent plant photosynthate into soil food webs regardless of their trophic complexity. Glob. Change Biol. 25, 3549–3561. 10.1111/gcb.14754

Cortet, J., Joffre, R., Elmholt, S., Krogh, P.H., 2003. Increasing species and trophic diversity of mesofauna affects fungal biomass, mesofauna community structure and organic matter decomposition processes. Biol. Fertil. Soils 37, 302–312. 10.1007/s00374-003-0597-2

Cotrufo, M.F., Wallenstein, M.D., Boot, C.M., Denef, K., Paul, E., 2013. The Microbial Efficiency-Matrix Stabilization (MEMS) framework integrates plant litter decomposition with soil organic matter stabilization: Do labile plant inputs form stable soil organic matter? Glob. Change Biol. 19, 988–995. 10.1111/gcb.12113

Crowther, T.W., Jones, T.H., Boddy, L., Baldrian, P., 2011. Invertebrate grazing determines enzyme production by basidiomycete fungi. Soil Biol Biochem 43, 2060–2068. 10.1016/j.soilbio.2011.06.003

de Groot, G.A., Jagers op Akkerhuis, G.A.J.M., Dimmers, W.J., Charrier, X., Faber, J.H., 2016. Biomass and Diversity of Soil Mite Functional Groups Respond to Extensification of Land Management, Potentially Affecting Soil Ecosystem Services. Front. Environ. Sci. 4. 10.3389/fenvs.2016.00015

de Mendiburu, F., 2020. Package “agricolae.” R Package Version 13–2.

de Vries, F.T., Caruso, T., 2016. Eating from the same plate? Revisiting the role of labile carbon inputs in the soil food web. Soil Biol Biochem 102, 4–9. 10.1016/j.soilbio.2016.06.023

de Vries, F.T., Shade, A., 2013. Controls on soil microbial community stability under climate change. Front. Microbiol. 4.

Edgar, R.C., Flyvbjerg, H., 2015. Error filtering, pair assembly and error correction for next-generation sequencing reads. Bioinformatics 31, 3476–3482. 10.1093/bioinformatics/btv401

Fanin, N., Kardol, P., Farrell, M., Nilsson, M.C., Gundale, M.J., Wardle, D.A., 2019. The ratio of Gram-positive to Gram-negative bacterial PLFA markers as an indicator of carbon availability in organic soils. Soil Biol. Biochem. 128, 111–114. 10.1016/j.soilbio.2018.10.010

Harkes, P., Elsen S. van den, Helder, H., 2025. Microbial DNA isolation from 1 gram soil.

Hawlena, D., Schmitz, O.J., 2010. Herbivore physiological response to predation risk and implications for ecosystem nutrient dynamics. Proc Natl Acad Sci 107, 15503–15507. 10.1073/pnas.1009300107

Hedlund, K., Griffiths, B., Christensen, S., Scheu, S., Setälä, H., Tscharntke, T., Verhoef, H., 2004. Trophic interactions in changing landscapes: responses of soil food webs. Basic Appl. Ecol. 5, 495–503. 10.1016/j.baae.2004.09.002

Hedlund, K., Öhrn, M.S., 2000. Tritrophic interactions in a soil community enhance decomposition rates. Oikos 88, 585–591. 10.1034/j.1600-0706.2000.880315.x

Hu, J., Gebremikael, M.T., Tytgat, B., Dumack, K., Hassi, U., Salehi Hosseini, P., Sleutel, S., Verleyen, E., De Neve, S., 2022. Combined selective gamma irradiation and pulverized soil inoculation for ecologically relevant soil microfauna studies. Appl. Soil Ecol. 169, 104223–104223. 10.1016/j.apsoil.2021.104223

Hunt, H.W., Coleman, D.C., Ingham, E.R., Ingham, R.E., Elliott, E.T., Moore, J.C., Rose, S.L., Reid, C.P.P., Morley, C.R., 1987. The detrital food web in a shortgrass prairie. Biol. Fertil. Soils 3, 57–68. 10.1007/BF00260580

Indorf, C., Dyckmans, J., Khan, K.S., Joergensen, R.G., 2011. Optimisation of amino sugar quantification by HPLC in soil and plant hydrolysates. Biol. Fertil. Soils 47, 387–396. 10.1007/s00374-011-0545-5

Joergensen, R.G., 2018. Amino sugars as specific indices for fungal and bacterial residues in soil. Biol. Fertil. Soils 54, 559–568. 10.1007/s00374-018-1288-3

Kallenbach, C.M., Frey, S.D., Grandy, A.S., 2016. Direct evidence for microbial-derived soil organic matter formation and its ecophysiological controls. Nat. Commun. 7, 1–10. 10.1038/ncomms13630

Kane, J.L., Kotcon, J.B., Freedman, Z.B., Morrissey, E.M., 2022. Fungivorous nematodes drive microbial diversity and carbon cycling in soil. Ecology e3844. 10.1002/ecy.3844

Laakso, J., Setälä, H., 1999. Population-and ecosystem-level effects of predation on microbial-feeding nematodes. Oecologia 120, 279–286.

Lang, B., Rall, B.C., Scheu, S., Brose, U., 2014. Effects of environmental warming and drought on size-structured soil food webs. Oikos 123, 1224–1233. 10.1111/j.1600-0706.2013.00894.x

Lejoly, J.D.M., Mason-Jones, K., Veen, G.F.C., 2026. A soil food web approach to integrate soil fauna into multitrophic biogeochemistry. Commun. Earth Environ. 7, 1–12. 10.1038/s43247-026-03322-4

Lenoir, L., Persson, T., Bengtsson, J., Wallander, H., Wirén, A., 2007. Bottom–up or top–down control in forest soil microcosms? Effects of soil fauna on fungal biomass and C/N mineralisation. Biol. Fertil. Soils 43, 281–294. 10.1007/s00374-006-0103-8

Lensing, J.R., Wise, D.H., 2006. Predicted climate change alters the indirect effect of predators on an ecosystem process. Proc. Natl. Acad. Sci. 103, 15502–15505. 10.1073/pnas.0607064103

Li, M.S.R., O’Rourke, D.R., Kaehler, B.D., Ziemski, M., Dillon, M.R., Foster, J.T., Bokulich, N.A., 2021. RESCRIPt: Reproducible sequence taxonomy reference database management. PLOS Comput. Biol. 17, e1009581. 10.1371/journal.pcbi.1009581

Li, Y., Veen, G.F., (Ciska)Hol, W.H.G., Vandenbrande, S., Hannula, S.E., ten Hooven, F.C., Li, Q., Liang, W., Bezemer, T.M., 2020. ‘Home’ and ‘away’ litter decomposition depends on the size fractions of the soil biotic community. Soil Biol. Biochem. 144, 107783–107783. 10.1016/j.soilbio.2020.107783

Liang, C., Amelung, W., Lehmann, J., Kästner, M., 2019. Quantitative assessment of microbial necromass contribution to soil organic matter. Glob. Change Biol. 25, 3578–3590. 10.1111/gcb.14781

Liang, C., Duncan, D.S., Balser, T.C., Tiedje, J.M., Jackson, R.D., 2013. Soil microbial residue storage linked to soil legacy under biofuel cropping systems in southern Wisconsin, USA. Soil Biol. Biochem. 57, 939–942. 10.1016/j.soilbio.2012.09.006

Liu, C., Cui, Y., Li, X., Yao, M., 2021. microeco: an R package for data mining in microbial community ecology. FEMS Microbiol. Ecol. 97, fiaa255. 10.1093/femsec/fiaa255

Lucas, J.M., McBride, S.G., Strickland, M.S., 2020. Trophic level mediates soil microbial community composition and function. Soil Biol. Biochem. 143, 1–10. 10.1016/j.soilbio.2020.107756

Mangiafico, S., 2021. Package “rcompanion.” R Package Version 2327.

Martikainen, E., Huhta, V., 1990. Interactions between nematodes and predatory mites in raw humus soil: a microcosm experiment. Rev. Ecol. Biol. Sol 27, 13–20.

Martinez-Almoyna, C., Saillard, A., Zinger, L., Lionnet, C., Arnoldi, C., Foulquier, A., Gielly, L., Piton, G., Münkemüller, T., Thuiller, W., 2022. Differential effects of soil trophic networks on microbial decomposition activity in mountain ecosystems. Soil Biol. Biochem. 172, 108771. 10.1016/j.soilbio.2022.108771

Mielke, L., Taubert, M., Cesarz, S., Ruess, L., Küsel, K., Gleixner, G., Lange, M., 2022. Nematode grazing increases the allocation of plant-derived carbon to soil bacteria and saprophytic fungi, and activates bacterial species of the rhizosphere. Pedobiol. -J. Soil Ecol. 90, 150787–150787. 10.1016/j.pedobi.2021.150787

Mikola, J., Setälä, H., 1998. No evidence of trophic cascades in an experimental microbial-based soil food web. Ecology 79, 153–164. 10.1890/0012-9658(1998)079[0153:NEOTCI]2.0.CO;2

Miyashita, T., Niwa, S., 2006. A test for top-down cascade in a detritus-based food web by litter-dwelling web spiders. Ecol. Res. 21, 611–615. 10.1007/s11284-006-0155-0

Nielsen, U.N., Ayres, E., Wall, D.H., Bardgett, R.D., 2011. Soil biodiversity and carbon cycling: a review and synthesis of studies examining diversity–function relationships. Eur. J. Soil Sci. 62, 105–116. 10.1111/j.1365-2389.2010.01314.x

Orwin, K.H., Dickie, I.A., Holdaway, R., Wood, J.R., 2018. A comparison of the ability of PLFA and 16S rRNA gene metabarcoding to resolve soil community change and predict ecosystem functions. Soil Biol Biochem 117, 27–35. 10.1016/j.soilbio.2017.10.036

Otto, S.B., Berlow, E.L., Rank, N.E., Smiley, J., Brose, U., 2008. Predator diversity and identity drive interaction strength and trophic cascades in a food web. Ecology 89, 134–144. 10.1890/07-0066.1

Parada, A.E., Needham, D.M., Fuhrman, J.A., 2016. Every base matters: Assessing small subunit rRNA primers for marine microbiomes with mock communities, time series and global field samples. Environ. Microbiol. 18, 1403–1414. 10.1111/1462-2920.13023

Pedregosa, F., Varoquaux, G., Gramfort, A., Michel, V., Thirion, B., Grisel, O., Blondel, M., Prettenhofer, P., Weiss, R., Dubourg, V., Vanderplas, J., Passos, A., Cournapeau, D., 2011. Scikit-learn: Machine Learning in Python. J. Mach. Learn. Res. 12, 2825–2830.

Postma-Blaauw, M.B., De Goede, R.G.M., Bloem, J., Faber, J.H., Brussaard, L., 2010. Soil biota community structure and abundance under agricultural intensification and extensification. Ecology 91, 460–473. 10.1890/09-0666.1

Prommer, J., Walker, T.W.N., Wanek, W., Braun, J., Zezula, D., Hu, Y., Hofhansl, F., Richter, A., 2020. Increased microbial growth, biomass, and turnover drive soil organic carbon accumulation at higher plant diversity. Glob. Change Biol. 26, 669–681. 10.1111/gcb.14777

Quast, C., Pruesse, E., Yilmaz, P., Gerken, J., Schweer, T., Yarza, P., Peplies, J., Glöckner, F.O., 2013. The SILVA ribosomal RNA gene database project: Improved data processing and web-based tools. Nucleic Acids Res. 41, 590–596. 10.1093/nar/gks1219

Quist, C.W., Gort, G., Mooijman, P., Brus, D.J., van den Elsen, S., Kostenko, O., Vervoort, M., Bakker, J., van der Putten, W.H., Helder, J., 2019. Spatial distribution of soil nematodes relates to soil organic matter and life strategy. Soil Biol. Biochem. 136, 107542. 10.1016/j.soilbio.2019.107542

Quist, C.W., van der Putten, W.H., Thakur, M.P., 2020. Soil predator loss alters aboveground stoichiometry in a native but not in a related range-expanding plant when exposed to periodic heat waves. Soil Biol Biochem 150. 10.1016/j.soilbio.2020.107999

R Core Team, 2023. R: The R project for statistical computing.

Ripple, W.J., Estes, J.A., Schmitz, O.J., Constant, V., Kaylor, M.J., Lenz, A., Motley, J.L., Self, K.E., Taylor, D.S., Wolf, C., 2016. What is a trophic cascade? Trends Ecol. Evol. 31, 842–849. 10.1016/j.tree.2016.08.010

Rizzuto, M., Leroux, S.J., Schmitz, O.J., 2024. Rewiring the carbon cycle: A theoretical framework for animal-driven ecosystem carbon sequestration. J. Geophys. Res. Biogeosciences 129, e2024JG008026. 10.1029/2024JG008026

Saetre, P., 1998. Decomposition, microbial community structure, and earthworm effects along a birch-spruce soil gradient. Ecology 79, 834–846. 10.2307/176583

Salas, E., Gorfer, M., Bandian, D., Wang, B., Kaiser, C., Wanek, W., 2023. A rapid and sensitive assay to quantify amino sugars, neutral sugars and uronic acid necromass biomarkers using pre-column derivatization, ultra-high-performance liquid chromatography and high-resolution mass spectrometry. Soil Biol. Biochem. 177, 108927. 10.1016/j.soilbio.2022.108927

Schmitz, O.J., Beckerman, A.P., O’Brien, K.M., 1997. Behaviorally mediated trophic cascades: Effects of predation risk on food web interactions. Ecology 78, 1388–1399. 10.1890/0012-9658(1997)078[1388:BMTCEO]2.0.CO;2

Schmitz, O.J., Krivan, V., Ovadia, O., 2004. Trophic cascades: the primacy of trait-mediated indirect interactions. Ecol. Lett. 7, 153–163. 10.1111/j.1461-0248.2003.00560.x

Schmitz, O.J., Wilmers, C.C., Leroux, S.J., Doughty, C.E., Atwood, T.B., Galetti, M., Davies, A.B., Goetz, S.J., 2018. Animals and the zoogeochemistry of the carbon cycle. Science 362, 1–10. 10.1126/science.aar3213

Sechi, V., De Goede, R.G.M., Rutgers, M., Brussaard, L., Mulder, C., 2018. Functional diversity in nematode communities across terrestrial ecosystems. Basic Appl. Ecol. 30, 76–86. 10.1016/j.baae.2018.05.004

Shurin, J.B., Borer, E.T., Seabloom, E.W., Anderson, K., Blanchette, C.A., Broitman, B., Cooper, S.D., Halpern, B.S., 2002. A cross-ecosystem comparison of the strength of trophic cascades. Ecol. Lett. 5, 785–791. 10.1046/j.1461-0248.2002.00381.x

Sinsabaugh, R.L., Lauber, C.L., Weintraub, M.N., Ahmed, B., Allison, S.D., Crenshaw, C., Contosta, A.R., Cusack, D., Frey, S., Gallo, M.E., Gartner, T.B., Hobbie, S.E., Holland, K., Keeler, B.L., Powers, J.S., Stursova, M., Takacs-Vesbach, C., Waldrop, M.P., Wallenstein, M.D., Zak, D.R., Zeglin, L.H., 2008. Stoichiometry of soil enzyme activity at global scale. Ecol. Lett. 11, 1252–1264. 10.1111/j.1461-0248.2008.01245.x

Sommer, N.R., Alshwairikh, Y.A., Arietta, A.Z.A., Skelly, D.K., Buchkowski, R.W., 2023. Prey metabolic responses to predators depend on predator hunting mode and prey antipredator defenses. Oikos e09664, e09664. 10.1111/oik.09664

Strickland, M.S., Hawlena, D., Reese, A., Bradford, M.A., Schmitz, O.J., 2013. Trophic cascade alters ecosystem carbon exchange. Proc. Natl. Acad. Sci. 110, 11035–11038. 10.1073/pnas.1305191110

Stromberger, M.E., Keith, A.M., Schmidt, O., 2012. Distinct microbial and faunal communities and translocated carbon in Lumbricus terrestris drilospheres. Soil Biol. Biochem. 46, 155–162. 10.1016/j.soilbio.2011.11.024

Thakur, M.P., Geisen, S., 2019. Trophic regulations of the soil microbiome. Trends Microbiol. 27, 771–780. 10.1016/j.tim.2019.04.008

Thakur, M.P., Herrmann, M., Steinauer, K., Rennoch, S., Cesarz, S., Eisenhauer, N., 2015. Cascading effects of belowground predators on plant communities are density-dependent. Ecol. Evol. 5, 4300–4314. 10.1002/ece3.1597

Trap, J., Bonkowski, M., Plassard, C., Villenave, C., Blanchart, E., 2016. Ecological importance of soil bacterivores for ecosystem functions. Plant Soil 398, 1–24. 10.1007/s11104-015-2671-6

Tsiafouli, M.A., Thébault, E., Sgardelis, S.P., de Ruiter, P.C., van der Putten, W.H., Birkhofer, K., Hemerik, L., de Vries, F.T., Bardgett, R.D., Brady, M.V., Bjornlund, L., Jørgensen, H.B., Christensen, S., Hertefeldt, T.D., Hotes, S., Gera Hol, W.H., Frouz, J., Liiri, M., Mortimer, S.R., Setälä, H., Tzanopoulos, J., Uteseny, K., Pižl, V., Stary, J., Wolters, V., Hedlund, K., 2015. Intensive agriculture reduces soil biodiversity across Europe. Glob. Change Biol. 21, 973–985. 10.1111/gcb.12752

Vance, E.D., Brookes, P.C., Jenkinson, D.S., 1987. An extraction method for measuring soil microbial biomass C. Soil Biol Biochem 19, 703–707. 10.1016/0038-0717(87)90052-6

Wang, M., Ruan, W., Kostenko, O., Carvalho, S., Hannula, S.E., Mulder, P.P.J., Bu, F., van der Putten, W.H., Bezemer, T.M., 2019. Removal of soil biota alters soil feedback effects on plant growth and defense chemistry. New Phytol. 221, 1478–1491. 10.1111/nph.15485

Wardle, D.A., Yeates, G.W., 1993. The dual importance of competition and predation as regulatory forces in terrestrial ecosystems: evidence from decomposer food-webs. Oecologia 93, 303–306. 10.1007/BF00317685

Wieder, W.R., Grandy, A.S., Kallenbach, C.M., Bonan, G.B., 2014. Integrating microbial physiology and physio-chemical principles in soils with the MIcrobial-MIneral Carbon Stabilization (MIMICS) model. Biogeosciences 11, 3899–3917. 10.5194/bg-11-3899-2014

Yeates, G.W., Bongers, T., De Goede, R.G.M., Freckman, D.W., Georgieva, S.S., 1993. Feeding habits in soil nematode families and genera—An outline for soil ecologists. J. Nematol. 25, 315–331.

Zaitsev, A.S., Birkhofer, K., Ekschmitt, K., Wolters, V., 2018. Belowground tritrophic food chain modulates soil respiration in grasslands. Pedosphere 28, 114–123. 10.1016/S1002-0160(18)60008-6

Zelles, L., 1999. Fatty acid patterns of phospholipids and lipopolysaccharides in the characterisation of microbial communities in soil: A review. Biol. Fertil. Soils 29, 111–129. 10.1007/s003740050533

