## Supplementary tables and figures for "Top predators in soil food webs increase carbon cycling efficiency"

*Suppl. Table 1. Initial conditions for experiment*

|  | Units | Average | Standard error |
| --- | --- | --- | --- |
| Microbial biomass carbon | μg.g-soil^-1^ | 13.6 | 3.5 |
| Microbial biomass nitrogen | μg.g-soil^-1^ | 1.9 | 0.2 |
| Leucine amino peptidase | nmol.g-soil^-1^.h^-1^ | 27.5 | 3.7 |
| Cellobiohydrolase | nmol.g-soil^-1^.h^-1^ | 128.1 | 9.9 |
| β-1,4-glucosidase | nmol.g-soil^-1^.h^-1^ | 433.2 | 34.5 |
| Acid phosphatase | nmol.g-soil^-1^.h^-1^ | 165.7 | 8.0 |
| β-1,4-N-acetyl-glucosaminidase | nmol.g-soil^-1^.h^-1^ | 166.7 | 10.6 |
| Arylsulfatase | nmol.g-soil^-1^.h^-1^ | 101.6 | 7.5 |
| β-1,4-xylosidase | nmol.g-soil^-1^.h^-1^ | 147.5 | 15.0 |
| Total extracellular enzymatic activity | nmol.g-soil^-1^.h^-1^ | 1170.3 | 61.6 |
| Muramic acid | μM.l^-1^ | 10.2 | 1.6 |
| Glucosamine + Mannosamine | μM.l^-1^ | 22.1 | 1.8 |
| Galactosamine | μM.l^-1^ | 47.0 | 3.3 |
| Total amino sugars | μM.l^-1^ | 79.3 | 6.1 |
| Gram positive bacteria | μg.g-soil^-1^ | 3.12 | 0.08 |
| Gram negative bacteria | μg.g-soil^-1^ | 2.71 | 0.07 |
| Fungi | μg.g-soil^-1^ | 1.05 | 0.2 |

*Suppl. Table 2. One-way analysis of variance of variables measured after one week (or Kruskal-Wallis test if indicated in parenthesis, when χ^2^  reported instead of F). Variables for which data was transformed with Tukey’s ladder of powers to ensure normality of residuals are depicted with an *.*

|  |  | Df | *F/χ^2^* | *p* |
| --- | --- | --- | --- | --- |
| Litter mass loss | Biota treatment | 3 | 3.45 | < .05 |
|  | Residuals | 28 |  |  |
| Microbial biomass carbon | Biota treatment | 3 | 0.44 | .728 |
|  | Residuals | 28 |  |  |
| Microbial biomass nitrogen* | Biota treatment | 3 | 0.93 | .438 |
|  | Residuals | 28 |  |  |
| Leucine amino peptidase* | Biota treatment | 3 | 3.30 | < .05 |
|  | Residuals | 28 |  |  |
| Cellobiohydrolase | Biota treatment | 3 | 1.61 | .209 |
|  | Residuals | 28 |  |  |
| β-1,4-glucosidase* | Biota treatment | 3 | 1.42 | .259 |
|  | Residuals | 28 |  |  |
| Acid phosphatase* | Biota treatment | 3 | 0.74 | .536 |
|  | Residuals | 28 |  |  |
| β-1,4-N-acetyl-glucosaminidase | Biota treatment | 3 | 0.83 | .491 |
|  | Residuals | 28 |  |  |
| Arylsulfatase | Biota treatment | 3 | 1.22 | .320 |
|  | Residuals | 28 |  |  |
| β-1,4-xylosidase* | Biota treatment | 3 | 1.63 | .204 |
|  | Residuals | 28 |  |  |
| Total extracellular enzyme activity | Biota treatment | 3 | 0.61 | .612 |
|  | Residuals | 28 |  |  |
| Gram-positive bacteria | Biota treatment | 3 | 5.41 | <.01 |
|  | Residuals | 28 |  |  |
| Gram-negative bacteria | Biota treatment | 3 | 2.22 | .108 |
|  | Residuals | 28 |  |  |
| Fungi | Biota treatment | 3 | 0.24 | .866 |
|  | Residuals | 28 |  |  |
| Total nematode abundance (Kruskal-Wallis) | Biota treatment | 1 | 0.10 | .753 |
| Bacterivorous nematode abundance (Kruskal-Wallis) | Biota treatment | 1 | 0.18 | .674 |
| Fungivorous nematode abundance* | Biota treatment | 1 | 0.27 | .615 |
|  | Residuals | 14 |  |  |
| Predatory nematode abundance | Biota treatment | 1 | 2.72 | .121 |
|  | Residuals | 14 |  |  |

*Suppl. Table 3. One-way analysis of variance of variables measured after two weeks (or Kruskal-Wallis test if indicated in parenthesis, when χ^2^  reported instead of F). Variables for which data was transformed with Tukey’s ladder of powers to ensure normality of residuals are depicted with an *.*

|  |  | Df | *F/χ^2^* | *p* |
| --- | --- | --- | --- | --- |
| Litter mass loss | Biota treatment | 3 | 1.08 | .372 |
|  | Residuals | 28 |  |  |
| Microbial biomass carbon | Biota treatment | 3 | 0.29 | .836 |
|  | Residuals | 28 |  |  |
| Microbial biomass nitrogen | Biota treatment | 3 | 1.47 | .243 |
|  | Residuals | 28 |  |  |
| Leucine amino peptidase* | Biota treatment | 3 | 0.51 | .682 |
|  | Residuals | 28 |  |  |
| Cellobiohydrolase* | Biota treatment | 3 | 1.73 | .183 |
|  | Residuals | 28 |  |  |
| β-1,4-glucosidase* | Biota treatment | 3 | 1.01 | .402 |
|  | Residuals | 28 |  |  |
| Acid phosphatase* | Biota treatment | 3 | 0.29 | .835 |
|  | Residuals | 28 |  |  |
| β-1,4-N-acetyl-glucosaminidase (Kruskal-Wallis) | Biota treatment | 3 | 1.11 | .774 |
| Arylsulfatase | Biota treatment | 3 | 0.76 | .527 |
|  | Residuals | 28 |  |  |
| β-1,4-xylosidase* | Biota treatment | 3 | 0.68 | .573 |
|  | Residuals | 28 |  |  |
| Total extracellular enzyme activity | Biota treatment | 3 | 1.27 | .304 |
|  | Residuals | 28 |  |  |
| Total nematode abundance | Biota treatment | 1 | 11.97 | <.01 |
|  | Residuals | 14 |  |  |
| Bacterivorous nematode abundance* | Biota treatment | 1 | 6.71 | <.05 |
|  | Residuals | 14 |  |  |
| Fungivorous nematode abundance | Biota treatment | 1 | 7.35 | <.05 |
|  | Residuals | 14 |  |  |
| Carnivorous nematode abundance | Biota treatment | 1 | 0.60 | .452 |
|  | Residuals | 14 |  |  |

*Suppl. Table 4. One-way analysis of variance of extracellular enzymatic activities and amino sugars after five weeks. Variables for which data was transformed with Tukey’s ladder of powers to ensure normality of residuals are depicted with an *.*

|  |  | Df | *F* | *p* |
| --- | --- | --- | --- | --- |
| Leucine amino peptidase* | Biota treatment | 3 | 3.73 | < .05 |
|  | Residuals | 28 |  |  |
| Cellobiohydrolase* | Biota treatment | 3 | 1.34 | .283 |
|  | Residuals | 28 |  |  |
| β-1,4-glucosidase* | Biota treatment | 3 | 2.24 | .105 |
|  | Residuals | 28 |  |  |
| Acid phosphatase* | Biota treatment | 3 | 0.88 | .464 |
|  | Residuals | 28 |  |  |
| β-1,4-N-acetyl-glucosaminidase | Biota treatment | 3 | 3.11 | <.05 |
|  | Residuals | 28 |  |  |
| Arylsulfatase | Biota treatment | 3 | 1.93 | .148 |
|  | Residuals | 28 |  |  |
| β-1,4-xylosidase | Biota treatment | 3 | 2.04 | .131 |
|  | Residuals | 28 |  |  |
| Total extracellular enzyme activity | Biota treatment | 3 | 2.43 | .086 |
|  | Residuals | 28 |  |  |
| Muramic acid* | Biota treatment | 3 | 1.87 | .160 |
|  | Residuals | 26 |  |  |
| Glucosamine + Mannosamine* | Biota treatment | 3 | 1.12 | .358 |
|  | Residuals | 26 |  |  |
| Galactosamine | Biota treatment | 3 | 2.93 | .052 |
|  | Residuals | 26 |  |  |
| Total amino sugars | Biota treatment | 3 | 1.31 | .294 |
|  | Residuals | 26 |  |  |
| Microbial biomass carbon | Biota treatment | 3 | 1.94 | .146 |
|  | Residuals | 28 |  |  |
| Microbial biomass nitrogen | Biota treatment | 3 | 1.33 | .283 |
|  | Residuals | 28 |  |  |
| Total nematode abundance | Biota treatment | 1 | 3.17 | .097 |
|  | Residuals | 14 |  |  |
| Bacterivorous nematode abundance | Biota treatment | 1 | 1.15 | .302 |
|  | Residuals | 14 |  |  |
| Nematode energy channel | Biota treatment | 1 | 5.48 | <.05 |
|  | Residuals | 14 |  |  |
| Microbial energy channel* | Biota treatment | 3 | 5.1 | <.01 |
|  | Residuals | 28 |  |  |

*Suppl. Table 5. Permutational analysis of variance using Bray-Curtis distance. For pairwise comparisons, the Bonferroni correction for multiple comparisons was applied to calculate the adjusted p value.*

|  | Df | Sum of squares | R^2^ | *F* | *p* |
| --- | --- | --- | --- | --- | --- |
| Model | 3 | 0.80 | 0.19 | 2.23 | < .001 |
| Residuals | 28 | 3.34 | 0.81 |  |  |
| Total | 31 | 4.14 | 1.00 |  |  |
| *Pairwise comparisons* | | | | | |
| Microbiome - Nematodes | | | 0.08 | 1.29 | .46 |
| Microbiome - Nematodes + Mites | | | 0.15 | 2.42 | < .01 |
| Microbiome - Mites | | | 0.10 | 1.62 | .05 |
| Nematodes - Nematodes + Mites | | | 0.21 | 3.83 | < .05 |
| Nematodes - Mites | | | 0.16 | 2.65 | < .01 |
| Nematodes + Mites - Mites | | | 0.10 | 1.64 | < .05 |

*Suppl. Table. 6. Analysis of variance for dominant bacterial phyla (> 5 % relative abundance).*

|  |  | Df | F value | *p* value |
| --- | --- | --- | --- | --- |
| Chloroflexi | Biota treatment | 3 | 1.5 | .23 |
|  | Residuals | 28 |  |  |
| Proteobacteria | Biota treatment | 3 | 7.1 | < .01 |
|  | Residuals | 28 |  |  |
| Actinobacteriota | Biota treatment | 3 | 6.2 | < .01 |
|  | Residuals | 28 |  |  |
| Bacteroidota | Biota treatment | 3 | 1.1 | .36 |
|  | Residuals | 28 |  |  |
| Firmicutes | Biota treatment | 3 | 1.1 | .35 |
|  | Residuals | 28 |  |  |

### Supplementary figures

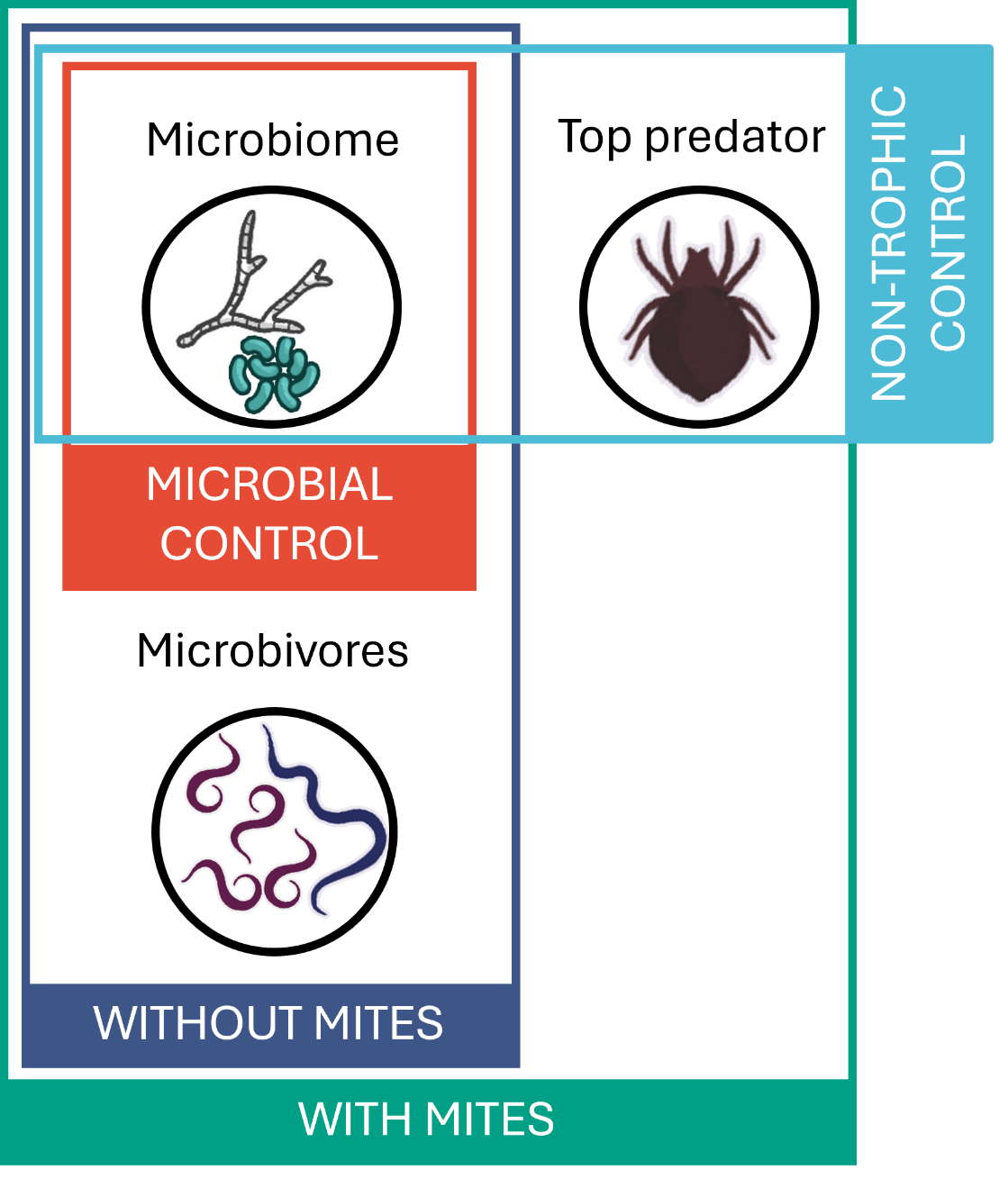

*Suppl. Fig. 1. Overview of the different soil biota treatments.*

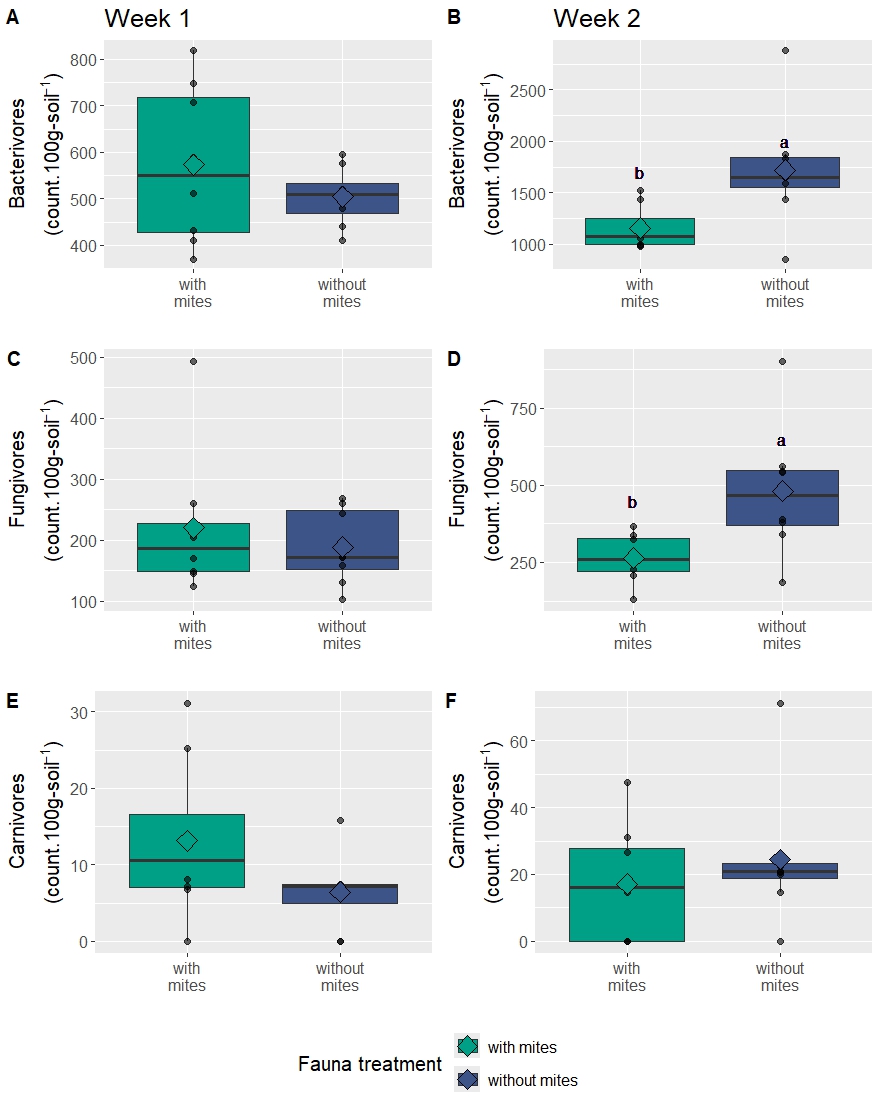

*Suppl. Fig. 2. Abundances of bacterivorous (A-B), fungivorous (C-D) and carnivorous nematodes after one and two weeks (n = 8) with different biota treatments (with mites: microbiome (M) + nematodes (N) + predatory mites (P), without mites: M + N, only treatments containing nematodes are displayed in the figure). Different letters indicate significant differences between treatments, according to the one-way analysis of variance (p < .05). Absence of letters means absence of significant differences (p > .05). Median and 25^th^ and 75^th^ percentiles (corresponding to the lower and upper hinges) as well as mean (diamond) and individual data points (dots).*

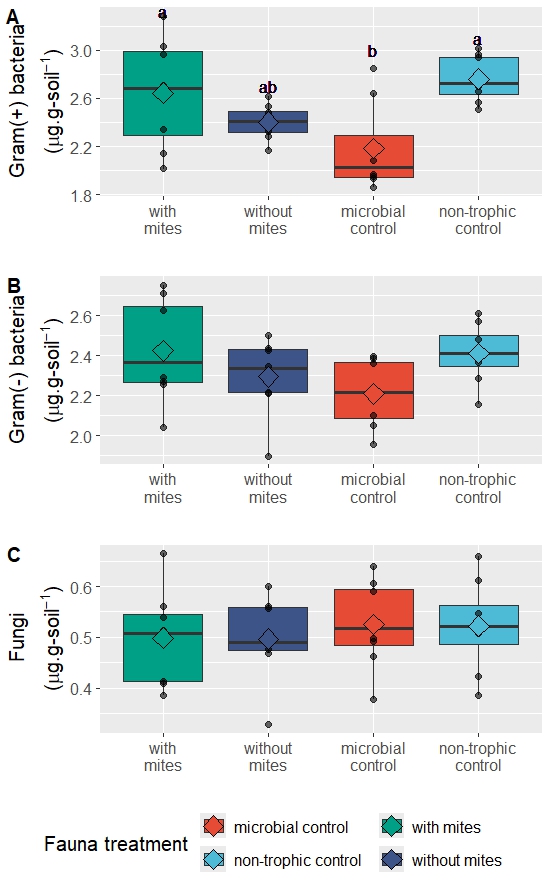

*Suppl. Fig. 3. Biomasses of Gram(+) (A) and Gram(-) (B) bacteria and fungi (C), estimated from phospholipid fatty acid analysis after one week (n = 8) with different biota treatments (with mites: microbiome (M) + nematodes (N) + predatory mites (P), without mites: M + N, microbial control: M, non-trophic control: M + P). Different letters indicate significant differences between treatments, according to the one-way analysis of variance (p < .05). Absence of letters means absence of significant differences (p > .05). Median and 25^th^ and 75^th^ percentiles (corresponding to the lower and upper hinges) as well as mean (diamond) and individual data points (dots).*

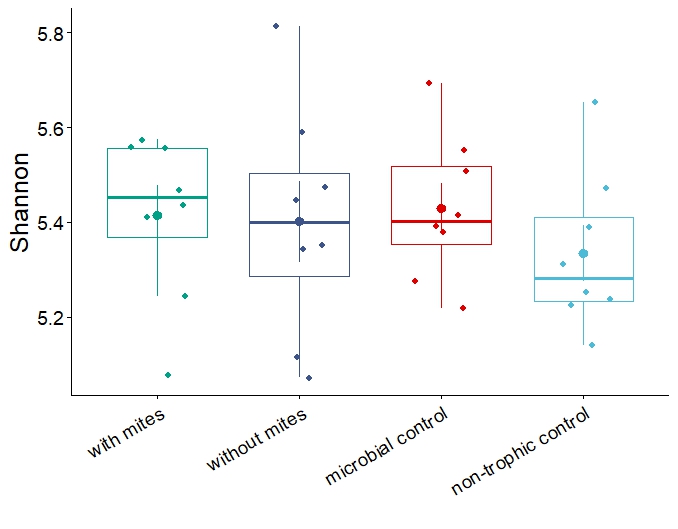

*Suppl. Fig. 4. Shannon diversity indices associated with each biota treatment (with mites: microbiome (M) + nematodes (N) + predatory mites (P), without mites: M + N, microbial control: M, non-trophic control: M + P) after five weeks (n=8). No significant differences were found between treatments, based on the results of the analysis of variance, as depicted by the same letters (p > .05). Median and 25^th^ and 75^th^ percentiles (corresponding to the lower and upper hinges) as well as mean (dot) and individual data points (diamonds).*

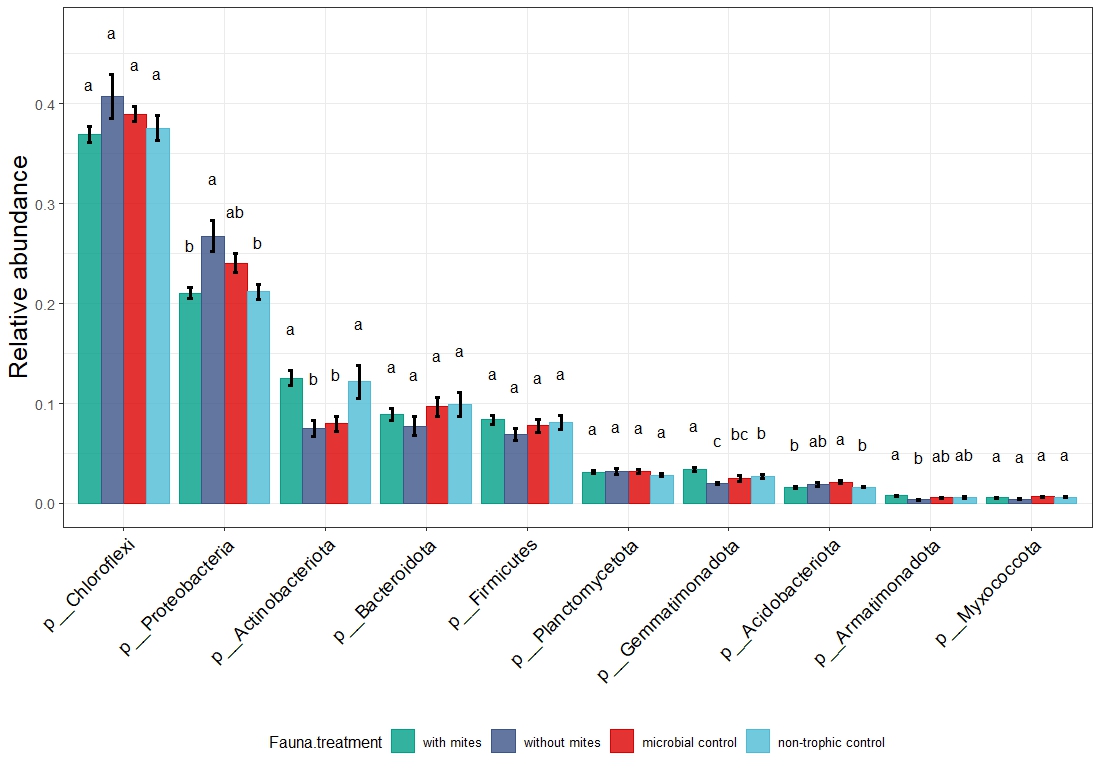

*Suppl. Fig. 5. Relative abundances of bacterial phyla obtained from 16S amplicon sequencing (n = 8) with error bars representing one standard error with different biota treatments (with mites: microbiome (M) + nematodes (N) + predatory mites (P), without mites: M + N, microbial control: M, non-trophic control: M + P). Different letters indicate significant differences between treatments, according to the one-way analysis of variance (p < .05). Absence of letters means absence of significant differences (p > .05). Statistical analyses were only conducted for phyla representing > 0.1 % of relative abundance.*

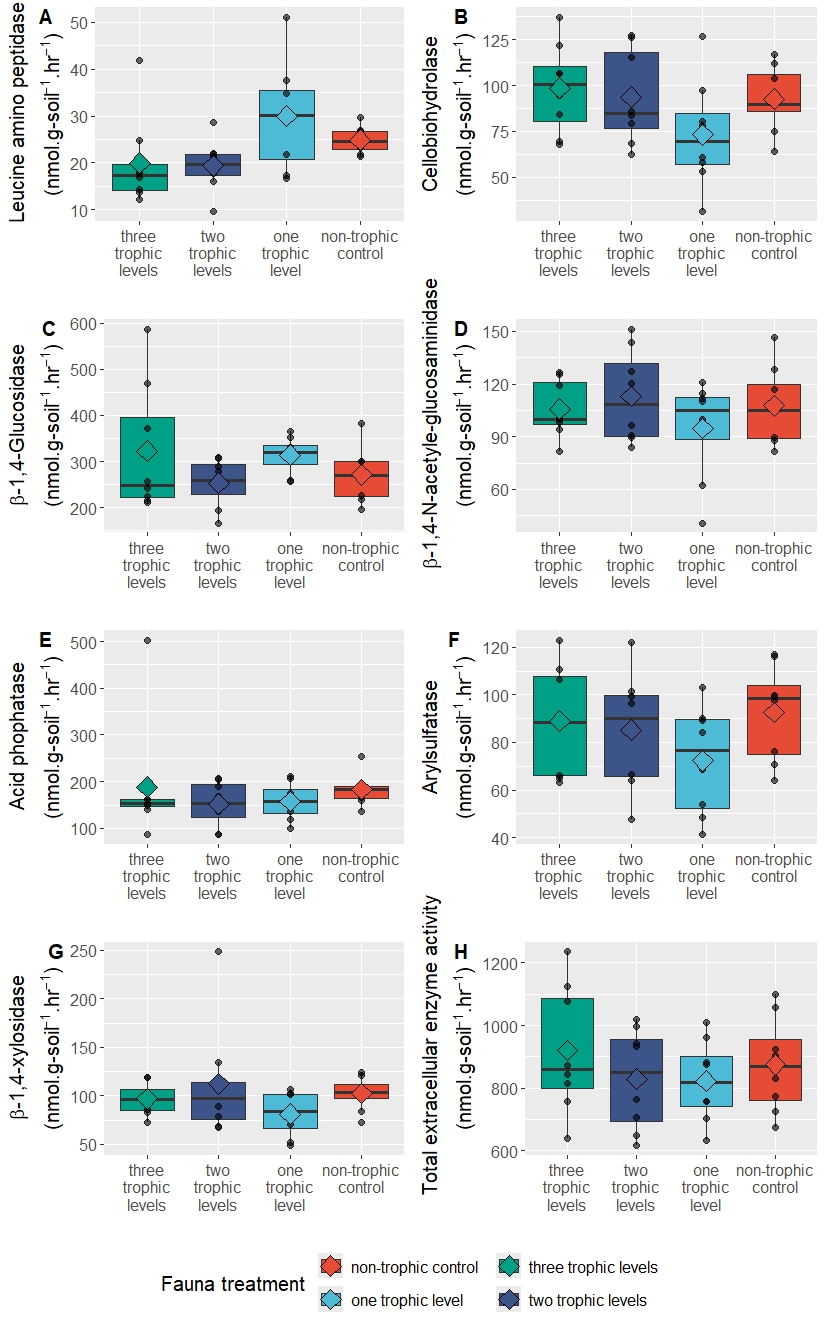

*Suppl. Fig. 6. Hydrolytic extracellular enzyme activities after one week (n = 8) with different biota treatments (with mites: microbiome (M) + nematodes (N) + predatory mites (P), without mites: M + N, microbial control: M, non-trophic control: M + P). Pairwise comparisons following one-way analysis of variance revealed no significant differences between treatments (p > .05). Median and 25^th^ and 75^th^ percentiles (corresponding to the lower and upper hinges) as well as mean (diamond) and individual data points (dots).*

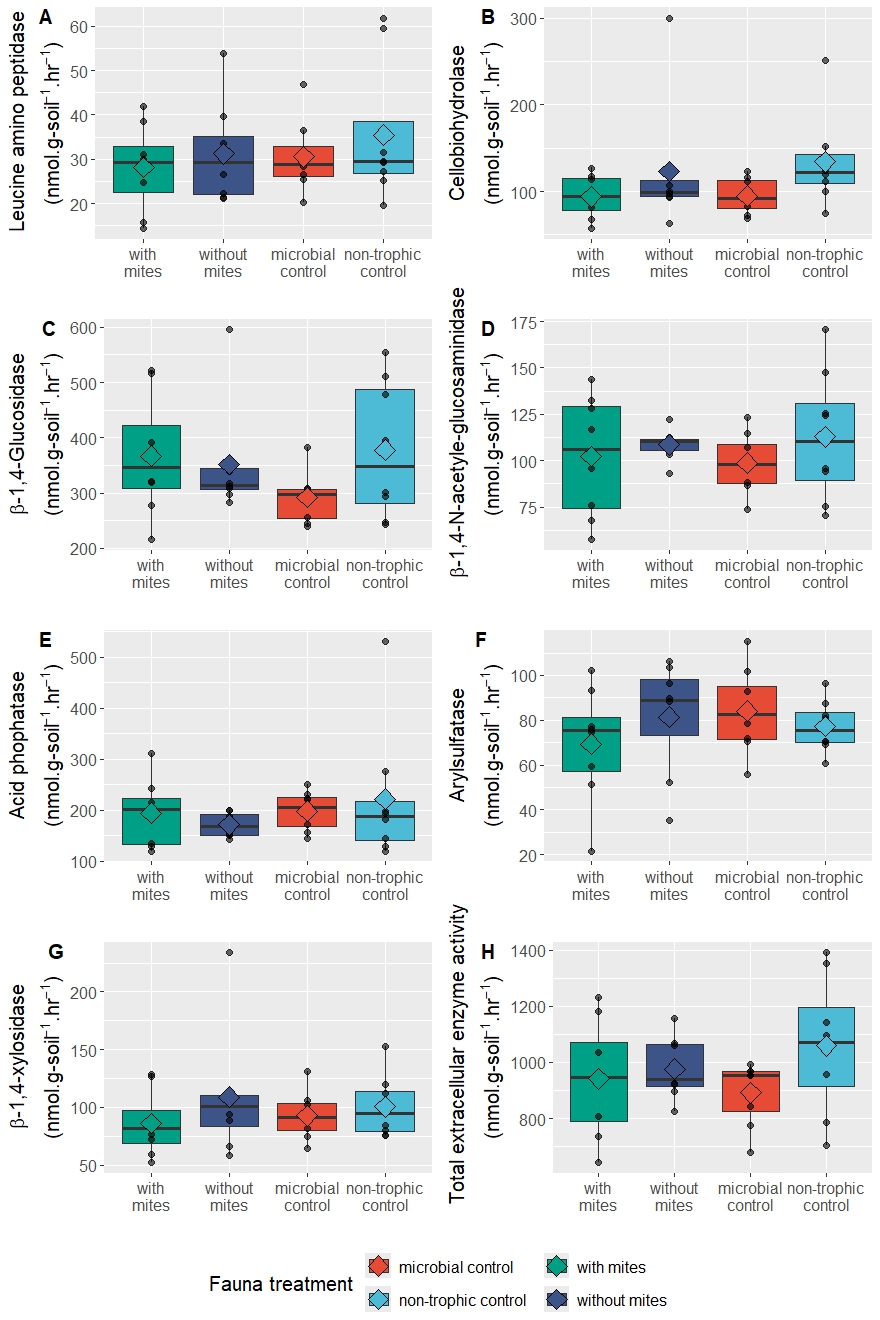

*Suppl. Fig. 7. Hydrolytic extracellular enzyme activities after two weeks (n = 8) with different biota treatments (with mites: microbiome (M) + nematodes (N) + predatory mites (P), without mites: M + N, microbial control: M, non-trophic control: M + P). Pairwise comparisons following one-way analysis of variance revealed no significant differences between treatments (p > .05). Median and 25^th^ and 75^th^ percentiles (corresponding to the lower and upper hinges) as well as mean (diamond) and individual data points (dots).*

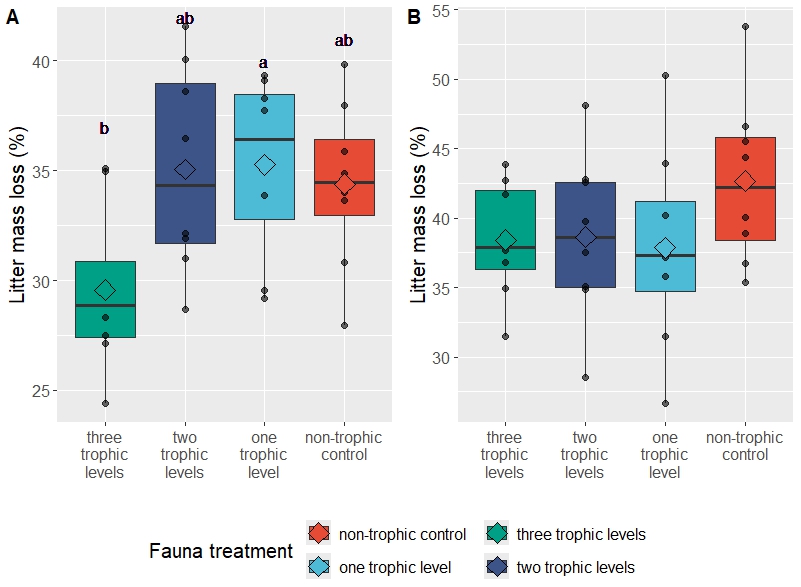

*Suppl. Fig. 8. Litter mass loss after one (A) and two (B) weeks (n = 8) with different biota treatments (with mites: microbiome (M) + nematodes (N) + predatory mites (P), without mites: M + N, microbial control: M, non-trophic control: M + P). Different letters indicate significant differences between treatments, according to the one-way analysis of variance (p < .05). Absence of letters means absence of significant differences (p > .05). Median and 25^th^ and 75^th^ percentiles (corresponding to the lower and upper hinges) as well as mean (diamond) and individual data points (dots).*

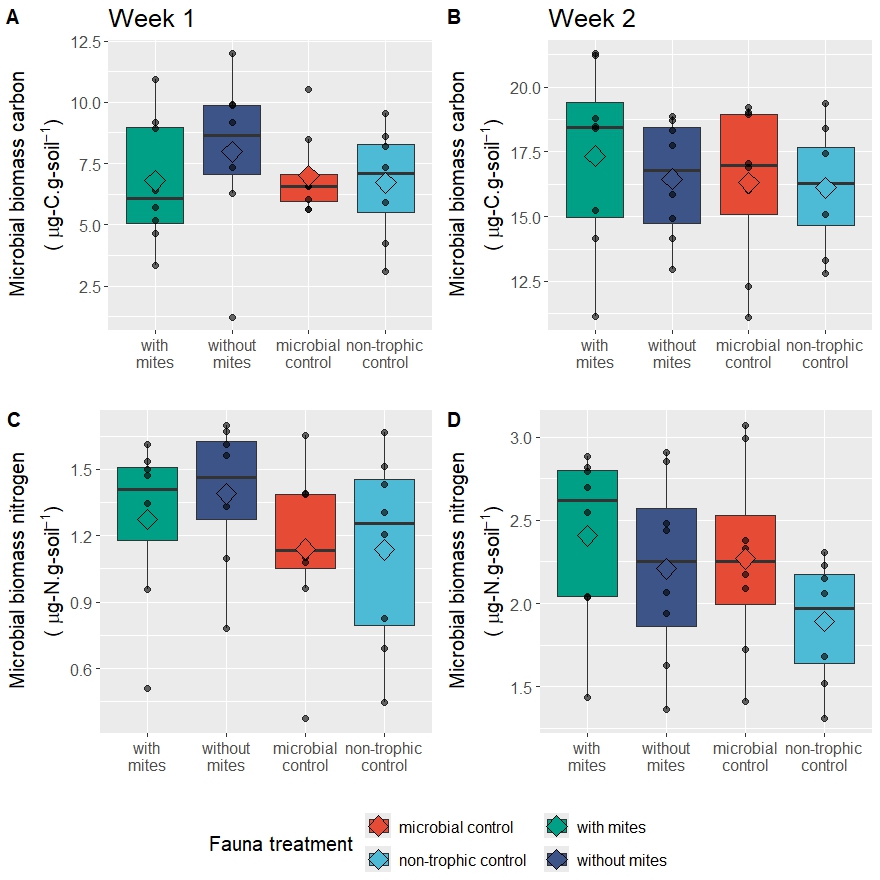

*Suppl. Fig. 9. Microbial biomass carbon (A-B) and nitrogen (C-D) after one and two weeks (n = 8) with different biota treatments (with mites: microbiome (M) + nematodes (N) + predatory mites (P), without mites: M + N, microbial control: M, non-trophic control: M + P). One-way analyses of variance revealed no significant differences between treatments (p > .05). Median and 25^th^ and 75^th^ percentiles (corresponding to the lower and upper hinges) as well as mean (diamond) and individual data points (dots).*
